# Ras/MAPK signalling intensity defines subclonal fitness in a mouse model of primary and metastatic hepatocellular carcinoma

**DOI:** 10.1101/2021.08.13.456223

**Authors:** Anthony Lozano, François-Régis Souche, Christel Ramirez, Serena Vegna, Guillaume Desandré, Anaïs Riviere, Valérie Dardalhon, Amal Zine El Aabidine, Philippe Fort, Leila Akkari, Urszula Hibner, Damien Grégoire

## Abstract

Quantitative differences in signal transduction are to date an understudied feature of tumour heterogeneity. The MAPK Erk pathway, which is activated in a large proportion of human tumours, is a prototypic example of distinct cell fates being driven by signal intensity. We have used primary hepatocyte precursors transformed with different dosages of an oncogenic form of Ras to model subclonal variations in MAPK signalling. Orthotopic allografts of Ras-transformed cells in immunocompromised mice gave rise to fast-growing aggressive tumours, both at the primary location and in the peritoneal cavity. Fluorescent labelling of cells expressing different oncogene levels, and consequently varying levels of MAPK Erk activation, highlighted the selection processes operating at the two sites of tumour growth. Indeed, significantly higher Ras expression was observed in primary as compared to metastatic tumours, despite the evolutionary trade-off of increased apoptotic death in the liver that correlated with high Ras dosage. Analysis of the immune tumoral microenvironment at the two locations suggests that fast metastatic growth in the immunocompromised setting is abrogated in immunocompetent animals due to efficient antigen presentation by peritoneal dendritic cells. Furthermore, our data indicate that, in contrast to the metastatic outgrowth, strong MAPK signalling is required in the primary liver tumours to resist elimination by NK cells. Overall, this study describes a quantitative aspect of tumour heterogeneity and highlights potential vulnerability of a subtype of hepatocellular carcinoma as a function of MAPK Erk signalling intensity.

## Introduction

Tumour subclonal heterogeneity is defined by cancer cells’ intrinsic properties and by phenotypic adaptations to extrinsic signals from the microenvironment (Marusyk et al., 2020). Specific tumour environments are shaped by the physiological constraints imposed by the tumour localisation and by the dynamic interactions with the growing tumour (Gerstung et al., 2020). The environment impacts the tumour phenotype, first by triggering adaptive responses from the highly plastic cancer cells (Pastushenko and Blanpain, 2019) and second by exerting selective pressures, leading to expansion of some, and elimination of other, tumoral subclones (Janiszewska et al., 2019). While the notion of inter- and intra-tumour heterogeneities and their consequences for personalized medicine have gained a strong momentum in the last decade (Losic et al., 2020; Molina-Sánchez et al., 2020), quantitative aspects leading to differences in the intensity of oncogenic signal transduction within the selected, clonal tumour cell populations (see e.g. (Gross et al., 2019)) are so far understudied.

Genomic analyses identified major oncogenic pathways for numerous tumour types. (Sanchez-Vega et al., 2018). For example, hepatocellular carcinoma (HCC) has been classified into several classes, defined by their genomic and physiopathological characteristics (Llovet et al., 2021). While some oncogenic driver mutations fall neatly in such defined categories, e.g. mutations that activate the β-catenin pathway or that inactivate the p53 tumour suppressor (Zucman-Rossi et al., 2015); others are frequently present in several different HCC types. An interesting example is the activation of the Ras/MAPK Erk pathway. Although mutations of Ras GTPases are found in only 2-4% of HCC patients, multiple activators and regulators of the pathway, such as FGF19, RSK2, RASAL1, RASSF1 and DUSP’s, are more frequently mutated. Altogether, the Ras/MAPK is estimated to be activated in 40 % of HCC (Delire and Stärkel, 2015; Lim et al., 2018).

Activation of Ras/MAPK Erk signalling is an essential feature of proliferating cells (Pagès et al., 1993) and the genetic abrogation of several of the components of the pathway gives rise to embryonic lethality due to a decrease in signal intensity (Dorard et al., 2017). Moreover, variations in signal intensity, duration and tissue-specific context give rise to distinct cellular outcomes (Lenormand et al., 1993; Marshall, 1995; Pouysségur et al., 2002).

In this work, we transformed hepatocyte precursors (BMEL, bipotent mouse embryonic liver cells) with different number of copies of an oncogenic mutant Ras (H-Ras^G12V^), which entailed variable intensity of signalling through the Ras/MAPK Erk pathway. We show that these cells tolerate a wide range of activated Ras dosage *in vitro*. In contrast, *in vivo* tumours display a much narrower range of active Ras levels and the consequent Erk signalling, likely attributable to selective pressures from the microenvironment. Specifically, cells that sustain low levels of Ras/MAPK Erk pathway activation are gradually eliminated from the liver primary tumours. Strikingly, they persist in secondary tumours that develop in the peritoneal cavity. Thus, specific microenvironments present in primary and in metastatic tumour locations, which differ in their macrophage, dendritic and NK cell immune landscape, select distinct tumoral subclones that are characterized by site-specific Ras signalling dosage.

## Results

### The phenotype of hepatic progenitors is affected by the Ras^G12V^ gene dosage

In order to investigate the consequences of distinct Ras oncogenic dosage on tumour development, we used BMEL, bi-potential precursors of the two epithelial hepatic lineages, derived from embryonic mouse liver (Akkari et al., 2010; Strick-Marchand and Weiss, 2002, 2003). BMEL are not transformed and retain major characteristics of primary hepatic progenitors, including the capacity to repopulate a damaged adult liver (Strick-Marchand et al., 2004). We have previously reported that the expression of an oncogenic form of Ras (H-Ras^G12V^) is sufficient to transform a subset of BMEL clones (Akkari et al., 2012; Bacevic et al., 2019), including the ones used for the current study. These cells express p19ARF and wild-type p53, but not p16-INK4A (Suppl Fig. 1), which could account for their escape from Ras-induced senescence (Xue et al., 2007).

Because cellular phenotypes are sensitive to the Ras oncogene dosage (Kerr et al., 2016; Mueller et al., 2018) and the signalling intensity of its downstream effector, the MAPK Erk pathway (Dikic et al., 1994; Traverse et al., 1994), we asked whether a specific level of Ras^G12V^ expression was required for hepatic tumour growth *in vivo*. To obtain cellular populations with distinct levels of Ras^G12V^ expression, BMEL cells were transduced with bi-cistronic lentiviral vectors encoding H-Ras^G12V^ and either a Venus or a mCherry fluorescent protein (Fig. 1A upper). Cells were sorted by flow cytometry on the basis of the fluorescence intensity, giving rise to populations that we named “Ras^LOW^” and “Ras^HIGH^” (Fig. 1A middle). Since the parental BMEL cells used in this study were clonally derived, the BMEL-Ras populations were isogenic, except for the copy number and insertion sites of the Ras^G12V^ - Venus/mCherry transgene and the consequent level of their expression. As expected, the expression levels of the fluorescent markers faithfully reflected the Ras oncogene mRNA levels (Fig. 1A lower). Importantly, the Ras^HIGH^ and Ras^LOW^ cells, while respectively enriched in strong or weak H-Ras^G12V^ expressors, were still heterogeneous with regard to Ras expression levels, the distribution of which presented an overlap between the two populations (Suppl Fig. 2). Neither the level of Ras expression nor the nature of the co-expressed fluorescent protein altered the *in vitro* proliferation rate of the transformed BMEL cells (Suppl Fig. 3).

**Figure 1:**
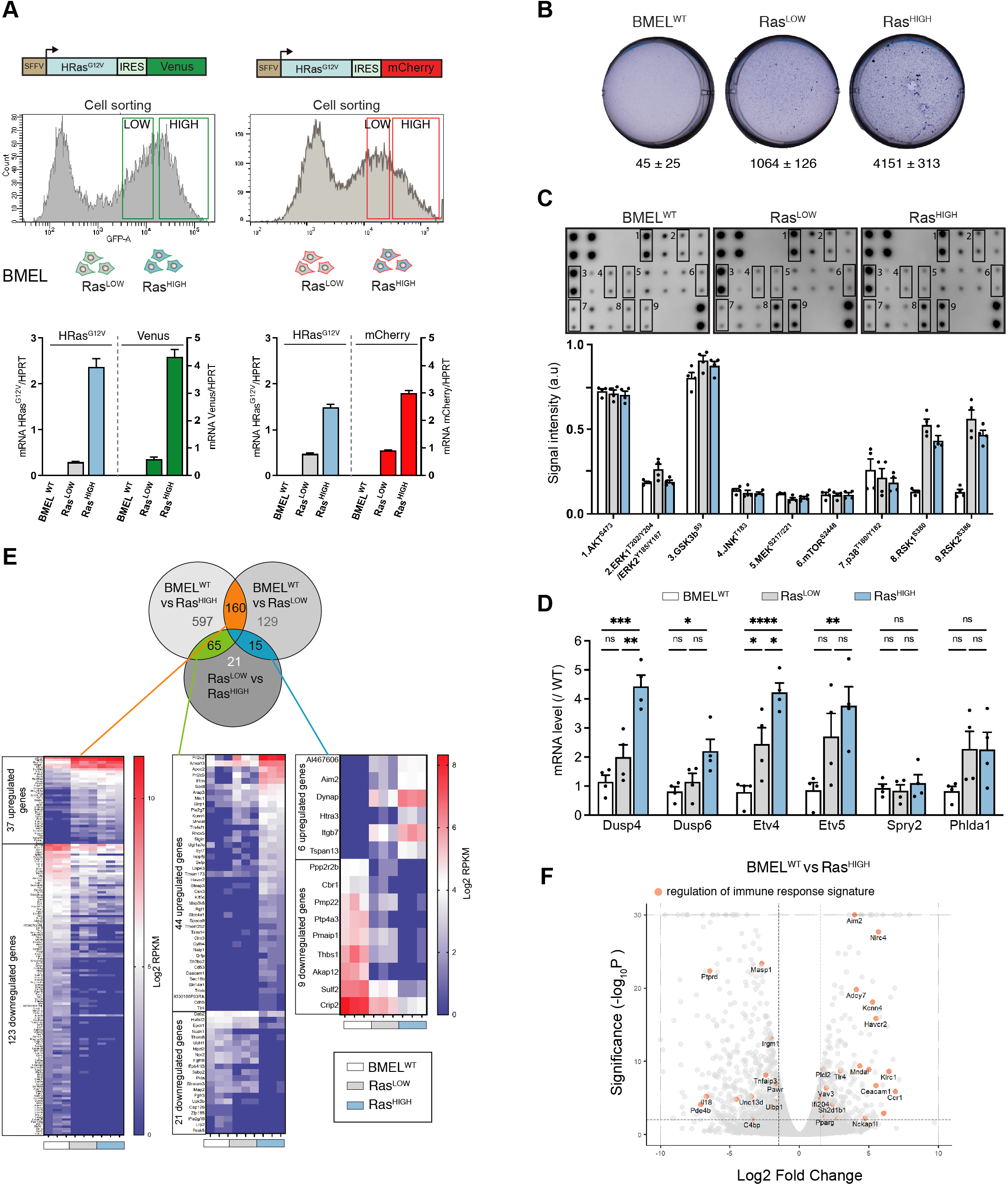
Ras^G12V^ oncogenic dosage controls the phenotype of BMEL cells. **A.** Bicistronic vectors (upper) were used to transduce BMEL cells that were sorted by flow cytometry according to fluorescence intensities (upper middle), giving rise to Ras^LOW^ and Ras^HIGH^ populations labelled with either Venus or mCherry (lower middle). Ras^G12V^, Venus and mCherry expression were quantified by RTqPCR (lower panels). **B.** Parental non-transformed BMEL («BMEL^WT^»), Ras^LOW^ and Ras^HIGH^ cells were seeded in soft agar, stained with crystal violet and counted after 21 days of culture. Representative wells and mean colony numbers +/- SEM from two independent experiments performed in triplicate are shown. **C.** Phosphoprotein array performed on BMEL^WT^, Ras^LOW^ and Ras^HIGH^ cell lines. Bottom panel shows the quantification of duplicates from two independent experiments, a.u: arbitrary units **D.** qPCR quantification of Erk target genes signature in Ras^LOW^ and Ras^HIGH^ BMEL cells normalized to expression in parental BMEL^WT^ cells. **E.** Venn diagram and heatmaps showing log2 RPKM values of genes regulated by Ras^G12V^ in BMEL cells detected by a RNASeq transcriptomic analysis (Log2FC >1.5, p-value<0.01). Genes expressed at low levels (all RPKM Log2 values <1) were removed from the analysis. From left to right: 160 genes were modulated by Ras similarly in Ras^LOW^ and Ras^HIGH^ populations, 65 genes were altered only in the Ras^HIGH^ cells and the expression of 15 genes was gradually modulated in the Ras^LOW^ and Ras^HIGH^ populations. **F.** Volcano plot presentation of deregulated genes in Ras^HIGH^ cells vs parental BMEL cells. Genes belonging to regulation of immune response signature (GO:0050776) are highlighted in orange. Ns not significant, *<0.05, **<0.01, ***<0.001 **** < 0.0001

While the parental BMEL cells did not form colonies when deprived of anchorage, Ras-transformed cells grew in soft agar, with the Ras^HIGH^ cells giving significantly more colonies than their Ras^LOW^ counterparts in this surrogate assay of transformation (Fig. 1B). Several major signal transduction pathways lay downstream of Ras activation. To gain insight into their activation status in H-Ras^G12V^-transformed BMEL cells, we assayed phosphorylation levels of a set of kinases (Fig. 1C). Oncogenic Ras had no effect on AKT^S473^ and the GSKb^S9^, two downstream targets of PI(3)K that were strongly phosphorylated both in the parental and in the Ras-transformed, cells. Similarly, no differences were detected for JNK and p38 pathway components. In contrast, although this assay did not reveal increased MEK or Erk activation, Ras signalling gave rise to a strong increase in the phosphorylation of Rsk 1 and 2, which are downstream targets of MAPK Erk signalling. Moreover, while the proteomic array assay was not sufficiently sensitive to distinguish between the Ras^LOW^ and Ras^HIGH^ signalling, the oncogenic dosage did translate into differences of the Ras/MAPK Erk signalling, as evidenced by distinct transcriptional signatures of known Erk delayed early target genes (Brant et al., 2017) (Fig. 1D).

In order to further investigate how the intensity of Ras signalling governs the hepatic transcriptional programmes, we performed RNAseq analysis of parental BMEL, Ras^LOW^ and Ras^HIGH^ cells in culture. Setting the thresholds at fold change log2 = 1.5 and p-value <0.01 identified over 1000 genes modulated by Ras^G12V^ signalling in these hepatic progenitor cells (Fig. 1E). The expression of one hundred sixty genes was modulated by Ras independently of its mean expression level in the tested population (Fig. 1E left, Suppl Table 1), thus representing a gene signature for the low threshold Ras/MAPK signalling. In a second group of 65 genes, target gene expression was modified only in the Ras^HIGH^ cells (Fig. 1E middle, Suppl Table 2). For these genes, a high intensity Ras signalling is required for the mRNA accumulation. Finally, the third group contains genes the expression of which correlated with the mean level of Ras expression. We identified 6 genes (*Al467606*, *Aim2*, *Dynap*, *Htra3*, *Itgb7*, *Tspan13*) whose expression was gradually increased in Ras^LOW^ and Ras^HIGH^ cell populations (Fig. 1E right, Suppl Table 3). Similarly, we found 9 genes (*Ppp2r2b*, *Cbr1*, *Pmp22*, *Ptp4a3*, *Pmaip1*, *Thbs1*, *Akap12*, *Sulf2*, *Crip2*) the expression of which was gradually repressed. Thus, our analysis identifies novel sets of quantitatively regulated Ras/MAPK target genes in hepatic cells. In the general framework of questioning the impact of Ras oncogenic dosage on the tumour-stroma interactions, we note the presence of genes in the GO category of immune response regulation (Fig. 1F).

### Specific Ras^G12V^ gene dosage is selected during tumour growth

Orthotopic injections of 10^5^ BMEL-Ras cells into immunocompromised recipients systematically gave rise within 3-4 weeks to hepatocellular tumours at the site of injection as well as to frequent extra-hepatic growth in the peritoneal cavity. First, we concentrated on primary tumours arising in the liver. Allografts of Ras^LOW^ and Ras^HIGH^ cells both produced poorly differentiated and fast growing, aggressive tumours, as witnessed by their overall size, high proliferation index and typically a multi-nodular morphology (Fig. 2A). In accordance with their superior capacity for anchorage independent growth (Fig. 1B), the Ras^HIGH^ cells gave rise to significantly larger tumours than their Ras^LOW^ counterparts, indicating that high level of Ras signalling conferred a selective advantage *in vivo*. To validate this observation, we next performed orthotopic injections of a 1:1 mix of Ras^HIGH^-Venus: Ras^LOW^-mCherry cells. The *in vivo* seeding efficiencies of Ras^HIGH^ and Ras^LOW^ cells were indistinguishable, since equal numbers of Venus and mCherry labelled cells were detected at early times (5 days) post-injection (Fig. 2B). However, as the tumours grew, a clear imbalance between the two populations became apparent, with Ras^HIGH^ cells becoming the dominant population in tumours already at day 10, which was further confirmed in full-grown tumours 21 days post-injection (Fig. 2B). This was not due to an artefactual effect of the fluorescent markers, as switching the fluorescent labels between the Ras^HIGH^ and Ras^LOW^ cells had no impact on the dominance of Ras^HIGH^ cells. Furthermore, cells labelled with Venus or mCherry that expressed comparable Ras levels contributed equally to orthotopic tumours (Fig. 2B lower).

**Figure 2:**
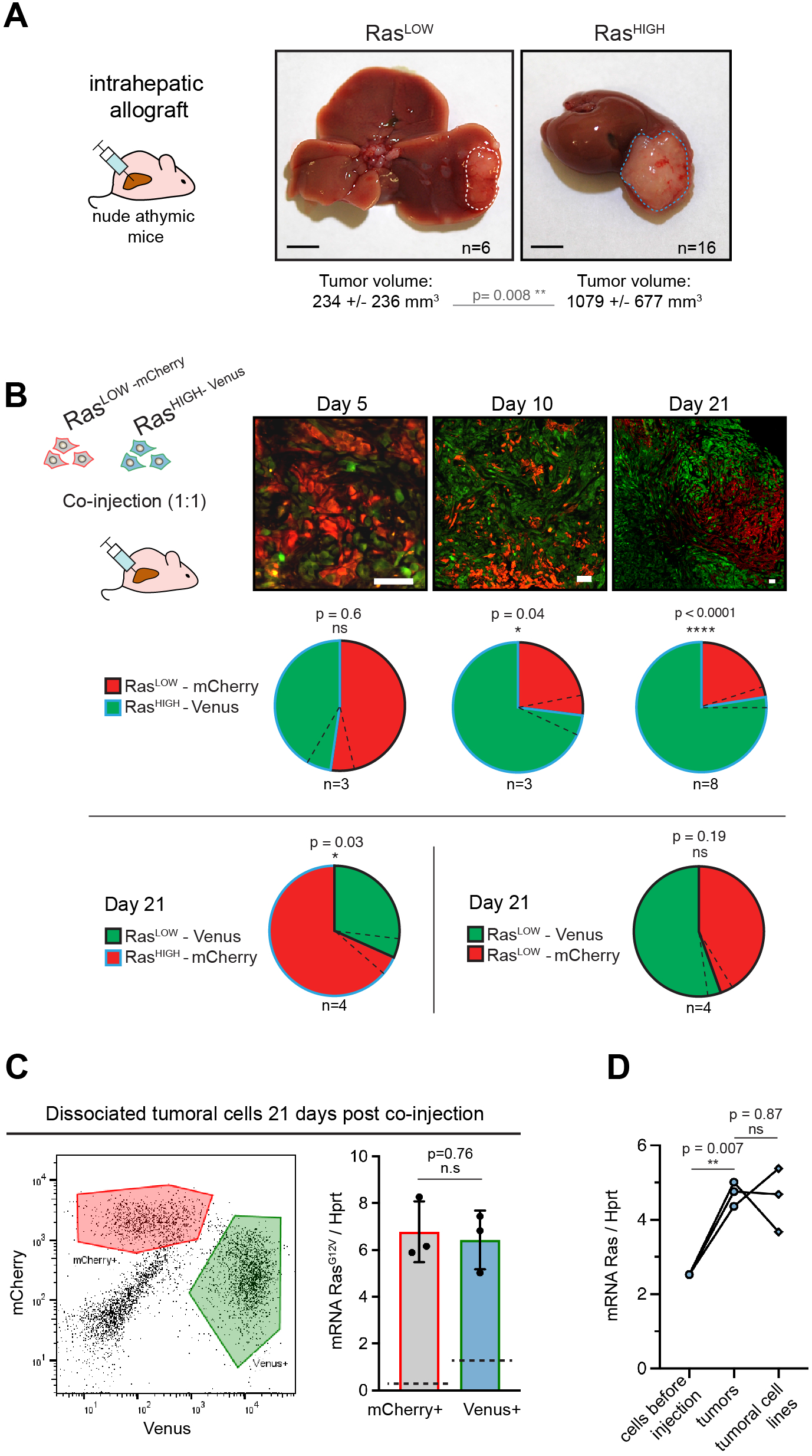
High level of Ras expression confers a selective advantage during in vivo tumour growth. **A.** Macros∞pic liver tumours three weeks after intrahepatic injection of 10^5^ Ras^LOW^ or Ras^HIGH^ BMEL cells, as indicated. Mean +/-S.D volumes are indicated. Scale bars: 5 mm **B.** Mice were injected orthotopically with a 1:1 mix of Ras^HIGH^-Venus (green) and Ras^LOW^-mCherry (red) cells and sacrificed at day 5,10 and 21, as indicated. Representative images of tumour sections are shown in the upper panels. Different magnifications were used to take into account the tumor growth (scale bars: 50 μm). Contribution of Ras^HIGH^ and Ras^LOW^ cells to the tumour composition was estimated by image analysis and flow cytometry, which gave identical results. Mean +/- SEM (dashed lines) are shown. The significance of deviations from the theoretical value of 50% (i.e no selective advantage) were calculated by the one sample t-test. **C.** Cells from dissociated tumours were sorted by flow cytometry (left panel) and the mean expression of Ras^G12V^ in Venus- and mCherry-labelled populations was quantified by RTqPCR. Mean +/- SEM from three independent tumors are shown. Dashed lines indicate the mean level of expression of Ras^G12V^ in the population of cells prior to injection **D.** Mean Ras^G12V^ expression level quantified by RTqPCR in Ras^HIGH^ cells before injection, cells freshly isolated from the tumours as well as in cell lines isolated from tumors kept in culture for 14 days. Ns not significant, *<0.05, **<0.01, ****<0.0001

We next enquired about the selective processes within each (i.e Ras^HIGH^ and Ras^LOW^) subpopulation within the tumours. Of note, and as expected from the gating in the cell sorting (Fig. 1A), while the mean Ras^G12V^ level was significantly different, there was an overlap of the oncogene dosage between the two injected populations (Suppl Fig. 2). Tumours collected at day 21 post-engraftment were dissociated, and Venus^+^ or mCherry^+^ cells were sorted by flow cytometry (Fig. 2C left panel). Interestingly, RTqPCR analysis specific for the Ras^G12V^ revealed that both cellular populations composing the tumour expressed identical level of the oncogene (Fig. 2C right panel). This indicates that selective pressures operating during the *in vivo* tumour growth allow the expansion of a population with a defined level of Ras oncogene expression, independently of which parental population they originate from. Since fewer cells in the Ras^LOW^ population fell within the range of this “optimal” oncogene dosage, they were underrepresented in the fully grown tumour at the end of the experiment. Finally, cell lines established from the orthotopic tumours maintained the high level of Ras expression *in vitro* (Fig. 2D), providing further support to the idea of pre-existing clones undergoing selection *in vivo*.

### Site specific tumour microenvironment selects distinct levels of oncogene dosage

Human hepatocellular carcinoma give rise mainly to intrahepatic metastases, but extrahepatic invasion also occurs (Katyal et al., 2000). This is also the case in our animal model of HCC orthotopic allografts. Indeed, in addition to intra-hepatic tumour spread, we consistently observed tumours in the peritoneal cavity, morphologically resembling the paired liver tumours (Fig. 3A). Extrahepatic tumours likely arose from cell leakage during the surgical procedure of intrahepatic injection, since peritoneal lavage performed after surgery contained low but detectable numbers (< 0.5%) of Venus-labelled cells. In this scenario, tumour initiation would be expected to be synchronous at the two locations.

**Figure 3:**
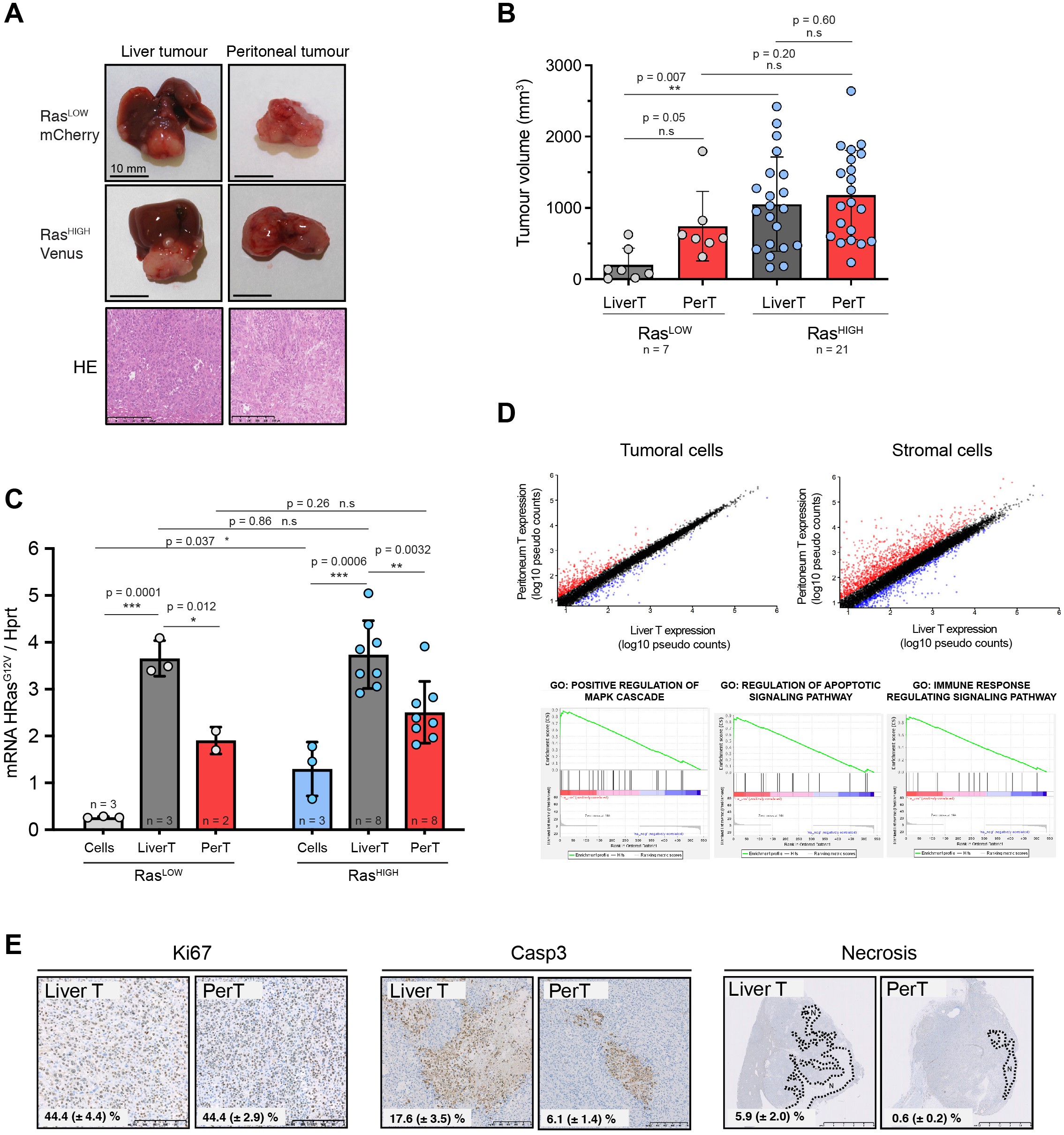
Distinct tumour environments select a site-specific range of optimal Ras signalling intensity. **A.** Representative images of paired liver and peritoneal tumours collected three weeks after injection of BMEL-Ras^LOW^ and Ras^HIGH^ cells, as indicated. Lower panel shows Hematoxylin/Eosin (HE) stainings. **B.** Quantification of liver and peritoneal tumour volumes from three independent experiments (mean +/- SD). LiverT: liver tumour, PerT: peritoneal tumour. Unpaired (*) (Ras^LOW^ vs Ras^HIGH^) or paired (#) (liverT vs perT) t-test p-values are indicated. **C.** Liver and peritoneal tumours resulting from injection of Ras^LOW^-mCherry or Ras^HIGH^-Venus cells, as indicated, were dissociated and fluorescent cells were sorted by flow cytometry. Ras^G12V^ expression levels in cells prior to injection and in cells purified from the tumours were quantified by Taqman RTqPCR. Mean +/- SD are shown. P-values were calculated by Student t-test: unpaired t-test (*<0.05, **<0.01, ***<0.001). **D.** Normalized enriched pathways from pre-ranked (Log2FC, Deseq) GSEA (GSEAv6: C5 Gene Ontology gen sets). Lower panels show the enrichment plot for the indicated GO terms: positive regulation of MAPK cascade, regulation of apoptotic signalling, immune response regulation. **E.** Immunohistochemical analysis of cell proliferation, apoptotic cell death and necrotic areas in Ras^HIGH^ liver and peritoneal tumours. Numbers on each panel represent mean+/-SEM from the analysis of duplicate tumour sections from 7 animals.

To gain further insight into the characteristics of tumour growth in the primary and metastatic sites, we analysed hepatic and peritoneal tumours generated following injections of either Ras^HIGH^ or Ras^LOW^ cell populations. Strikingly, whereas Ras^LOW^ cells gave significantly smaller hepatic tumours than Ras^HIGH^ cells, this difference was abolished with regard to peritoneal tumours (Fig. 3B). Altogether, these results confirm that the Ras^LOW^ population contained fewer cells capable of initiating and/or sustaining tumour growth in the liver. In contrast, there appears to be no selective advantage for Ras^HIGH^ cells to establish tumours in the peritoneal cavity.

To further investigate the hypothesis that tumorigenesis at both tissue locations was regulated by the intensity of Ras signalling, we improved the precision of the quantification of the *in vivo* oncogenic dosage. To do so, we used flow cytometry to separate tumoral from stromal cells derived from both liver primary tumours and from the matched peritoneal metastatic ones. RTqPCR analyses confirmed the increased oncogenic dosage in the *in vivo*-grown cells compared to the *in vitro*-grown parental ones and revealed a significantly higher mean level of Ras^G12V^ expression in the primary versus the metastatic tumours (Fig. 3C). Strikingly, the mean level of the oncogene expression was again independent of the origin of the engrafted cells (i.e. Ras^HIGH^ versus Ras^LOW^ populations). Overall, these data support the conclusion that tissue-specific selective pressures favour distinct oncogenic Ras levels.

To address the mechanistic basis for this phenomenon, we performed transcriptomic profiling of Venus+ tumour cells isolated from the primary and the metastatic sites. This analysis identified approximately 160 genes differentially expressed in the liver and peritoneal tumour cells (log2 fold change >1; p<0.05. Suppl. Table 4). Gene ontology analysis using GSEA identified 27 enriched gene sets (25 in liver and 2 in peritoneum; p-value < 0.01 and FDR < 0.1. Suppl. Table 5). Importantly, it confirmed the increase of MAPK signalling in the liver and identified apoptotic processes and the regulation of the immune response among the gene sets differentially regulated in the two locations (Fig. 3D). As expected, comparison of transcriptomic profiles from stromal cells of liver and peritoneal tumors pinpointed higher diversity (2260 genes for log2 fold change >1; p<0.05; Fig. 3D).

Both the hepatic and the peritoneal tumours were highly proliferative (Fig. 3E). Interestingly, while the cultured BMEL cells expressing either high or low levels of the Ras^G12V^ oncogene have comparable, low, apoptotic indexes, hepatic tumours contain a higher proportion of cleaved caspase 3 positive cells than their peritoneal counterparts (Fig. 3E). Moreover, in the liver, and much less so in the peritoneum, caspase-3 positive cells often surround large necrotic areas, indicating that the cell death in an expanding liver tumour is quite substantial, suggesting an evolutionary trade-off for the high oncogene dosage in the liver tumours.

### Immune cell contexture differs in primary and metastatic tumours

We next interrogated the interplay between the tumour cells and their local microenvironments that shape the oncogenic expression profile. The transcriptomic profiling of the stromal component of liver and peritoneal tumours revealed a number of differentially expressed genes, as expected for distinct tumour locations. Cibersortx deconvolution of the immune component of the stroma (Newman et al., 2019) did not show strong differences in the numbers of intra-tumoral macrophages at the two locations (Fig. 4A). However, independent RTqPCR profiling of macrophage polarization markers did suggest a more immunosuppressive, less inflammatory microenvironment in the peritoneum (Fig.4B, Suppl Fig. 4).

**Figure 4:**
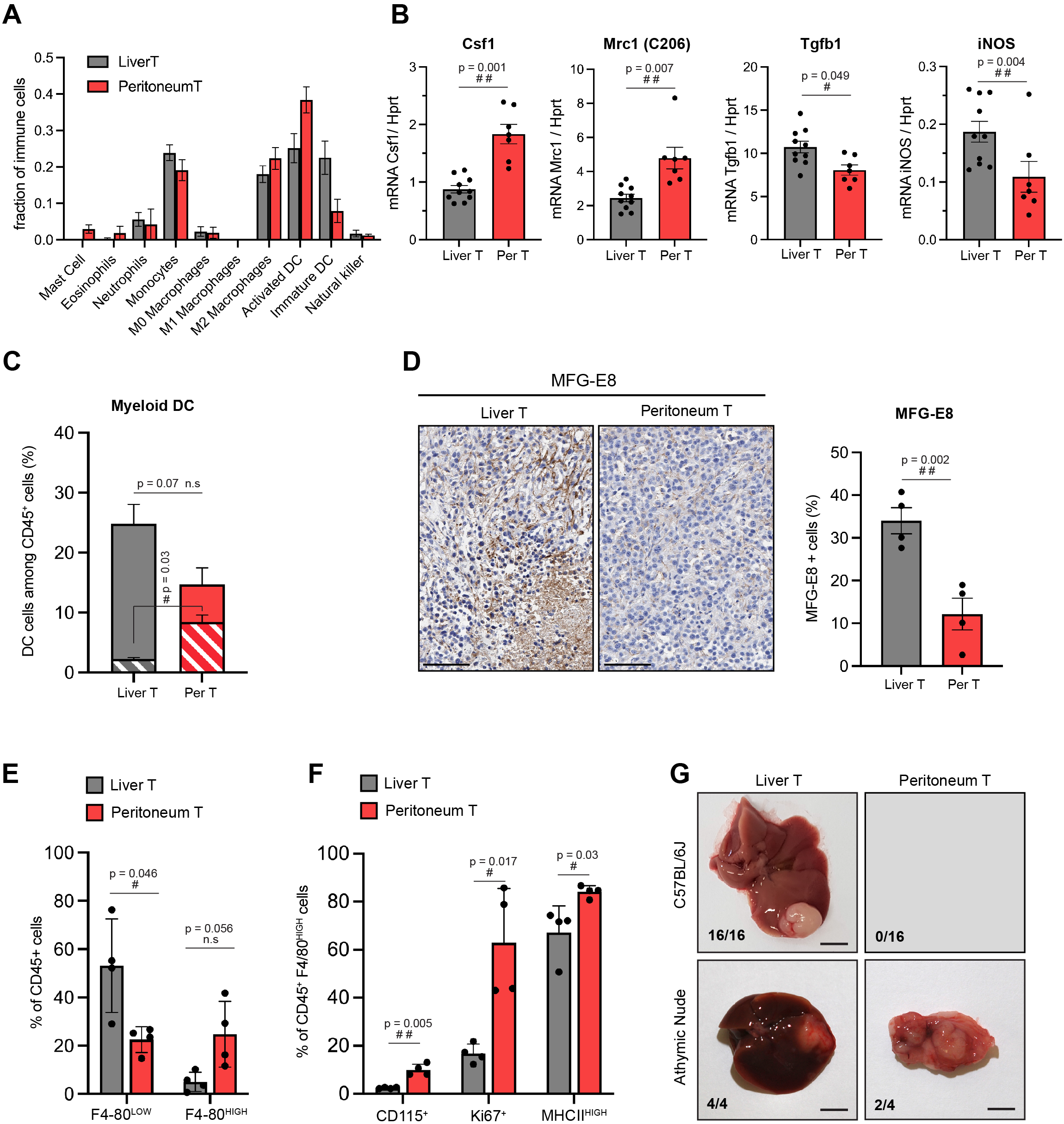
Site-specific immune contexts of primary and metastatic tumour locations. **A.** Cibersortx algorithm was used to perform cell type deconvolution of RNAseq data from stroma of matched liver and peritoneal tumours (n=4). Ten immune cell populations were included in the signature, as indicated. **B.** Macrophage polarization marker expression were analysed by RT-qPCR performed on RNA isolated from stromal cells sorted from liver and peritoneal tumours. **C.** Dendritic cells (DC) (CD45^+^, CD11c^+^, MHCII^high^, F4/80^-^, CD24^+^) from liver and peritoneal tumours were analysed by flow cytometry. The histogram represents the percentage of dendritic cells among the total CD45^+^ population, indicating the proliferating (Ki67^+^, dashed bars) and non-proliferating (Ki67^-^, filled bars) DC cells. **D.** Immunohistochemistry performed on liver and peritoneal tumour sections using anti MFG-E8 antibody, quantification of positive staining is shown in the right panel. Scale bars: 100 μm **E.** Flow cytometry analysis of CD45^+^CD11 b*Ly6G^-^Ly6C^-^ F4/80^|OW^ (monocyte-derived macrophages) and CD45^+^CD11 b*Ly6G^-^Ly6C^-^ F4/80^high^ (tissue-resident macrophages) from liver and peritoneal tumours. **F.** Quantification of CD115^+^, Ki67^+^ and MHCII^high^ cells among CD45^+^CD11b^+^Ly6GLy6CF4/80^high^ macrophages. H. 5000 cells (NRAS^G12D^ p53^KO^) were injected orthotopically in C57BL/6J mice and athymic nude mice, as indicated. Macroscopic images of representative liver and peritoneal tumours collected 3 weeks after injections (scale bar = 5mm). Numbers indicate the fraction of inoculated animals that developed tumours. All quantitative analyses were subjected to a paired student t-test. P-values: #<0.05, ##<0.01, ###<0.001

Interestingly, the presence of immunosuppressive tumour-associated macrophages (TAMs) did not deter from other antigen presenting cells activity in the peritoneal tumour microenvironment (TME). Indeed, our data suggested that a larger proportion of mature, activated dendritic cells (DC), as compared to immature ones, were present in this organ (Fig. 4A). Flow cytometry analyses confirmed that while the total number of DC (defined as CD45^+^ CD11b^+^ CD11c^+^ MHCII^high^ CD24^+^F4/80^-^) was not significantly different between the two TME, these cells showed an increase in proliferation in the peritoneum (Fig. 4C). Furthermore, the peritoneal DCs contained lower levels of the Milk-fact globule-EGFVIII (MFG-E8), a factor known to limit DC immunogenic potential (Fig. 4D) (Baghdadi et al., 2012; Peng and Elkon, 2012).

The phenotypes of macrophages in matched liver and peritoneal tumours also showed some interesting differences. The content of peripherally recruited monocyte-derived macrophages presenting low expression of F4/80, defined as CD45^+^CD11b^+^Ly6G^-^Ly6C^-^F4/80^low^, was significantly enriched in liver tumours (Fig. 4E). On the other hand, the differentiated, tissue resident-like, F4/80^high^ macrophages, showed a tendency for enrichment in the peritoneal HCC. These results suggest that macrophage subsets are distinct in the liver and peritoneal TME, with tissue-resident, F4/80^high^ cells, which are part of organ homeostasis maintenance (Dou et al., 2020) dominating the peritoneal TME while abundant F4/80^low^ immature TAMs infiltrate the liver. Interestingly, F4/80^high^ peritoneal macrophages displayed a superior proliferative capacity (Ki67+) and activation phenotype (MHCII+, CD115+) compared to liver macrophages (Fig. 4F). Coherent with our findings that more activated and less immature DC are present in peritoneal TME (Fig. 4A, D), these results suggest that the liver TME may represent a more immune tolerant environment for Ras^HIGH^ tumour cell outgrowth compared to the peritoneum, in which the activation levels of the Ras oncogene do not affect the dynamics of tumorigenesis. Thus, through both flow cytometry and RNAseq deconvolution analyses, our results highlight the differences in stromal composition of the tumours in the two locations, showing the complex interplay of innate immune cells that may participate in shaping the clonal selection in the tumour.

We reasoned that a likely consequence of increased numbers of activated DCs would be an improved efficiency of antigen presentation, thereby allowing a more efficient tumour clearance in the peritoneum. While such phenotype is not relevant in immunodeficient mice that are unable to mount a T-cell response, it should be detectable in immunocompetent animals. In order to test this prediction, we compared the outcomes of orthotopic injections of tumour cells in immunocompetent C57BL/6 and immunodeficient nude mice. For these autologous allografts, we took advantage of a C57BL/6 primary cell line that we derived from a tumour established by a hydrodynamic gene transfer of constitutively active Ras and a simultaneous CrispR/Cas9-mediated inactivation of the p53 tumour suppressor (Bacevic et al., 2019). Orthotopic allografts of 5000 cells performed in parallel on immunocompetent and immunodeficient mice gave rise to comparable macroscopic liver tumours within 3 weeks (Fig. 4G). However, in contrast to nude mice, of which 50% developed macroscopic peritoneal tumours, no metastatic growth was detected in immunocompetent recipients. These results are consistent with the hypothesis that a more effective antigen presentation in the peritoneum limits metastatic tumour growth in immunocompetent animals.

### Ras signalling modulates tumour cells interactions with stromal cells

We next questioned the mechanistic bases of the tumour and stroma crosstalk that might shape the quantitative differences in the oncogenic pathway activation. To do so, we looked for genes whose expression is specifically induced by high-intensity Ras signalling, both in the parental BMEL Ras^HIGH^ cells and in the liver, as compared, respectively, to the Ras^LOW^ cells and the peritoneal tumours (Fig. 5A). Out of four genes (*Ceacam1*, *Csn3*, *Selp*, *Tmem252*) that corresponded to this criterion, we focused our attention on CEACAM1 (Carcinoembryonic antigen-related cell adhesion molecule 1), a gene widely expressed in many cancer types and whose expression level has been reported to correlate with tumour progression (Dankner et al., 2017). It is also expressed on the surface of several immune cell subsets, including macrophages, T lymphocytes and NK cells, where homophilic interactions with the CEACAM1^+^ tumour cells abrogate NK-mediated cytotoxicity (Helfrich and Singer, 2019). In order to confirm that the observed Ras-mediated transcriptional activation translates into CEACAM1 expression on the surface of cancer cells, we performed FACS analyses on cells dissociated from the liver and peritoneal tumours. As expected, the majority of the non-hematopoietic (CD45^neg^) cells in both tumoral locations expressed the Venus fluorescent protein and were therefore the Ras^G12V^ expressing tumoral cells. CEACAM1 expressing cells were highly enriched in this population in the liver as compared to peritoneal tumours, corresponding respectively to 30.1 +/- 2.4 % (mean+/-SEM) and 14.2 +/- 3% (mean+/- SEM) of CD45^neg^ cells (Fig. 5B). Strikingly, the mean fluorescent intensities of the CEACAM1 labelling in the Venus+ populations were indistinguishable between the tumours at the two locations. This is consistent with the transcriptomic data indicating that high level of Ras signalling is needed to activate CEACAM1 gene expression in BMEL cells (Fig. 1E and 1F) and confirms that the cells that have the required Ras dosage are more abundant in the liver than in the peritoneal tumours.

**Figure 5:**
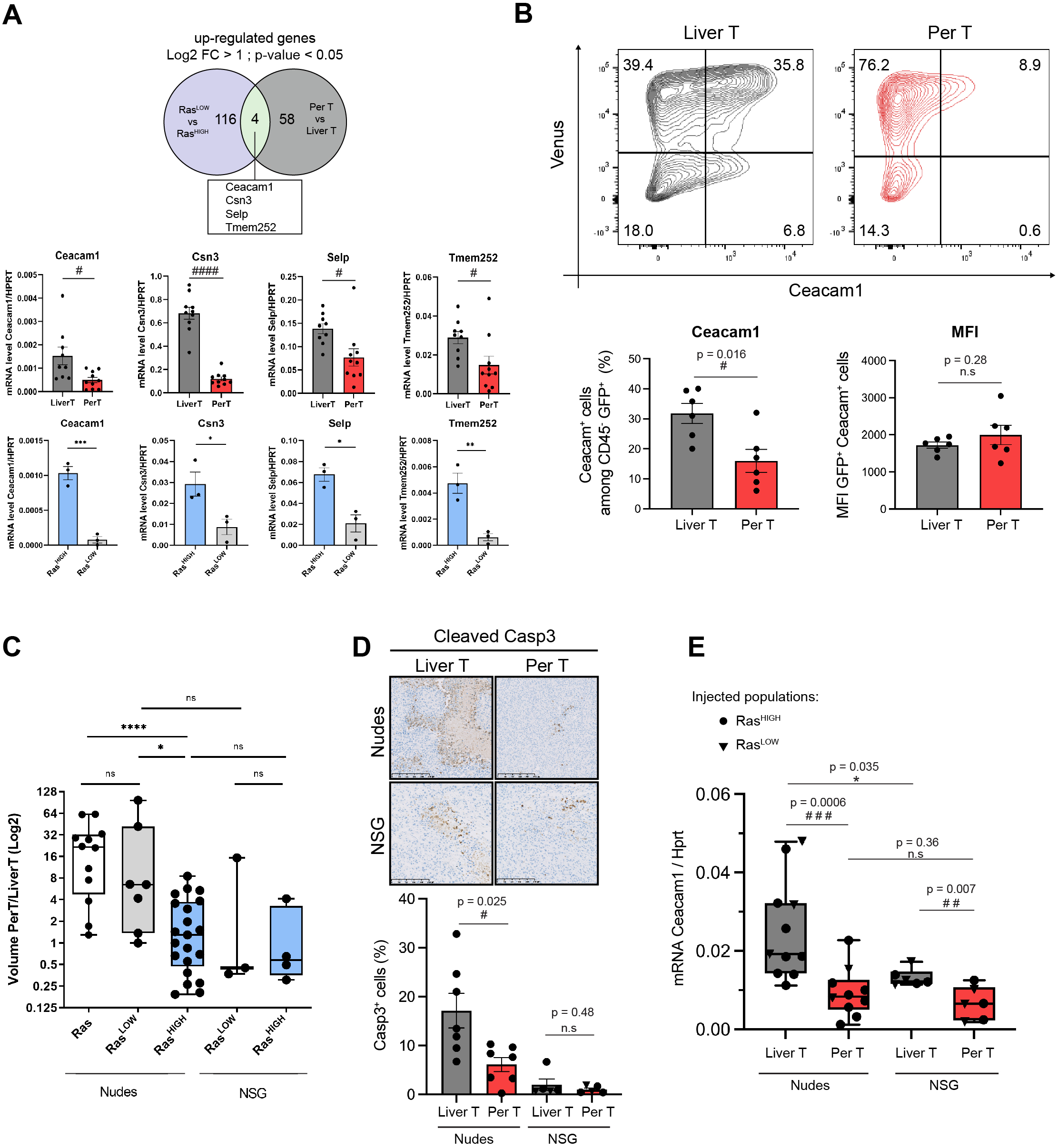
Contribution of natural killer cells to clonal selection in hepatic tumours. **A.** Venn diagram showing numbers of differentially expressed genes (Log2FC >1, p-value<0.05) from RNASeq data of liver vs peritoneal sorted tumoral cells and Ras^HIGH^ vs Ras^LOW^ BMEL cells. Bottom panel: RTqPCR quantification of the 4 genes upregulated both in the hepatic vs peritoneal tumours and in Ras^HIGH^ vs Ras^LOW^ cells. Upper panels: analysis of FACS sorted cells isolated from tumours, lower panels: analysis of in vitro grown cells. **B.** Flow cytometry analysis of CD45^-^ Venus+ Ceacam1+ cells isolated from the liver and the peritoneal tumours. Numbers indicate percentage of cells in each quadrant. Bottom panels show quantification (left) and mean fluorescent intensity (MFI, right) of the Ceacam?-ex-pressing tumour cells. **C.** Volume ratios of the matched peritoneal and liver tumors (PerT/LiverT) originating from injections of different cell populations, as indicated (non-sorted HRas^G12V^, Ras^LOW^, Ras^HIGH^) inoculated into athymic nude or NOD-SCID-gamma (NSG) mice. **D.** Immunohistochemical analysis of activated caspase 3 performed on tumours arising from Ras^LOW^ and Ras^HIGH^ cells injected in nude and NSG mice. The histogram shows the quantification of cleaved casp3 positive cells in randomly chosen fields. **E.** RTqPCR quantification of Ceacaml expression in sorted tumour cells from athymic nude and NSG mice. Unpaired Student t-test, p-values: ns not significant, *<0.05, **<0.01, ***<0.001 and ****<0.0001 p-values. Paired Student t-test, p-values: ns not significant, #<0.05, ##<0.01, ###<0.001.

Interestingly, while difference in the NK cell content in the primary and metastatic tumours did not reach statistical significance in the deconvolution of stromal cells RNAseq (Fig. 4A), immunofluorescence staining revealed distinct distribution of these cells: whereas NK infiltrate the hepatic tumours, they apparently tend to remain in the periphery of the peritoneal ones (Suppl Fig. 5 lower panels). To pinpoint the suspected involvement of NK cells in the selection of Ras^HIGH^/CEACAM^HIGH^ cells in liver tumours, we next analysed tumorigenesis in NOD scid gamma (NSG) mice. These animals are deeply immunosuppressed: in addition to lacking mature B and T cells they are notably devoid of NK cells (Yu et al., 2008). The tumour volumes were comparable in the NSG and in the nude mice. As in the nude mice, the NSG background was permissive for peritoneal metastatic growth, however, the majority of the animals displayed comparable tumour volumes at the two locations (Fig. 5C). Consistently, the number of caspase 3 positive cells was low for both liver and peritoneal tumours in NSG animals (Fig. 5D). These results support the idea that the discrimination against the Ras^LOW^ cells, both in the peritoneum and in the liver is alleviated in the NSG background. In support of this contention, compared to the nude mice, the hepatic tumours in the NSG background tolerated lower levels of CEACAM1 expression, while there was no significant difference for the peritoneal tumours between the two genetic backgrounds. Of note, however, the difference between CEACAM1 expression in the hepatic and peritoneal tumours was diminished, but not abolished in the NSG animals (Fig. 5E). Altogether, these results suggest that CEACAM1-driven NK inhibition participates in the clonal selection, but is not the sole element of the advantage afforded by the high oncogene dosage.

## Discussion

Multiple environmental and cell-autonomous mechanisms, organized in negative and positive feedback loops, cooperate to allow an exquisite regulation of the intensity and duration of signalling, the variation of which can give rise to specific and distinct cell fates. The Ras/MAPK Erk pathway is a prototypic example of such complex regulatory circuits (Murphy and Blenis, 2006), whose output is also relevant in the context of tumorigenesis. This has been shown for example in response to the loss of anchorage, where strong Erk signal is protective, while a moderate one is required for anoïkis (Zugasti et al., 2001). However, excessive Erk activation can be detrimental to the cell, provoking cell cycle arrest or senescence (Serrano et al., 1997; Woods et al., 1997). More recent single cell functional analyses further support the evidence for cell-autonomous sensitivity to signal intensity, but also reveal the heterogeneity of cellular response, both *in vitro* and *in vivo* (Gross et al., 2019; Kumagai et al., 2015; Molina-Sánchez et al., 2020).

Our understanding of how the availability of extracellular stimuli, or the deregulation of their sensors, translate into cellular responses is still incomplete. For example, although only 5% of the small GTPase Ras needs to be activated to give rise to full activation of Erk (Hallberg et al., 1994), a clear dosage effect has been reported in pancreatic and lung tumours for activating Ras mutations (East et al., 2021; Kerr et al., 2016; Mueller et al., 2018). Moreover, amplifications of the wild type allele of Ras that give rise to an allelic imbalance and the consequent modulation of the activity of the oncogenic Ras mutants result in distinct cellular phenotypes (Ambrogio et al., 2018; Bremner and Balmain, 1990). This further argues for the physiological importance of precise Ras dosage in a growing tumour.

Of note, whereas activating Ras mutations are rare in HCC (Zucman-Rossi et al., 2015), alterations in the intensity of MAPK Erk signalling participate in shaping the tumoral phenotype, as revealed by inactivating mutations of Rsk2, which is both an ERK target and a negative regulator of the pathway (Chan et al., 2021). Moreover, a recent analysis of transcriptional signatures classified HCC among tumours with significant Ras pathway activation phenotype (East et al., 2021). Because the same report highlighted the predictive value of strong versus weak Ras-driven transcriptional signature, it is relevant to model these phenotypes in HCC by controlled dosage of the Ras oncogene, as we have done in this study.

Our exploration of signal intensity that is optimal for tumour growth in the context of its specific microenvironment led to a description of a novel feature of tumour heterogeneity, manifested by quantitative dosage differences of oncogenic signalling in tumours developing at primary and metastatic locations. We note that the peritoneal tumours in our animal model likely do not result from a *bona fide* metastatic process. Nevertheless, they do represent tumoral growth at an extra-hepatic site and are thus a convenient model to analyse primary and metastatic-like tumour growth within the same animal.

Although several oncogenic signalling pathways lie downstream of Ras (Sanchez-Vega et al., 2018), in our experimental model Ras^G12V^ dosage correlates with the MAP kinase Erk activity, but not with the phosphoinositide 3-kinase. Whether such uncoupling is a general feature of HCC or simply a reflection of a strong PI(3)K/Akt signaling present in the non-transformed BMEL parental cells remains to be established.

Comparing tumours formed by BMEL-Ras cells expressing a wide range of Ras^G12V^ oncogene (“parental Ras” cells), with those initiated by cells with the narrower range of signalling intensities (Ras^HIGH^ and Ras^LOW^ cells) suggests the existence of a tradeoff in the tumours’ evolutionary history. Indeed, we have shown that cells expressing high oncogene levels are globally favoured in the primary liver tumours, however, their selective advantage is curtailed by a higher probability of undergoing programmed cell death. While the propensity to activate the senescence and/or apoptosis in response to oncogenic Ras signalling is well known (Serrano et al., 1997; Woods et al., 1997), the observation of diminished cell death in the tumours growing in NSG animals points to an additional, non-cell autonomous, mechanism that drives this response.

The immune response to cancer comprises a complex interplay of innate and adaptive components of immunity (Fridman et al., 2012; Hou et al., 2020; Nguyen et al., 2021). Concentrating our study on animals devoid of an adaptive immune response gave us access to interactions between the tumour and its innate immune component. Our results point to key differences in myeloid cell population subsets in the liver and peritoneum of tumour-bearing mice. Notably, the content of tumour-infiltrating, immunosuppressive macrophages, whose function in promoting liver carcinogenesis is well recognized (Dou et al., 2020; Ringelhan et al., 2018), was enhanced in Ras^HIGH^ liver tumours and correlated with an increased fraction of immature DCs when compared to the peritoneal TME. These results suggest that rather than total numbers of each myeloid populations, it is the phenotype of these cells, driven by the local cues, that regulates their ability to further engage an anti-tumour immunity and control tumour outgrowth. In line with this observation, recent studies have shown that identical oncogenic drivers have strikingly different consequences on the content and activation of TME cells in lung or pancreatic adenocarcinomas (Kortlever et al., 2017; Sodir et al., 2020). Hence, in addition to the tumour cells’ genetic landscape itself (Wellenstein and de Visser, 2018), the downstream activation of signalling nodes pertaining to dosage of a specific oncogene should be considered as a tumour-intrinsic feature shaping the TME, as recently also reported for pancreatic cancer (Ischenko et al., 2021).

It is of particular interest to consider the role played by natural killer cells as part of the organismal defence against HCC. NK activities are strongly dependent on, as yet incompletely understood, organ-specific features (Shi et al., 2011). Resident NK cells acquire cytolytic functions mediated through TNFα and granzyme B during hepatotropic viral infections, cirrhotic pathology and HCC (Sällberg and Pasetto, 2020). The viral adaptation to the NK-mediated attack includes the suppression of their function through the CEACAM1-mediated blockade of their activity (Suda et al., 2018). Our results are fully consistent with the idea that a similar mechanism of immune escape operates in primary tumours of HCC. In contrast to the liver, the peritoneal resident cells resemble immature splenic NK cells (Gonzaga et al., 2011), which may account for a lesser involvement of these cells in anti-tumour immunity in this location, as suggested by our data.

How can the organ-specific differences in primary and metastatic tumour growth be exploited in designing novel therapeutic approaches remains an open question. The observation that only a narrow range signalling intensity through a signal transduction pathway is compatible with tumour development raises hopes for the therapeutic efficacy of its modulation, as recently reported for stratification of HCC response to chemotherapeutic intervention based on the inactivating Rsk2 mutations that modulate the strength of the MAPK signalling (Chan et al., 2021). Similarly, only lung adenocarcinoma with a strong Ras activating signature respond to inhibitors of the MAPK Erk pathway (East et al., 2021). It remains to be seen if the Ras pathway activation signature proves to be useful for HCC stratification for defining clinically relevant treatment options.

## Materials & Methods

### BMEL preparation and cell culture

BMEL cells were isolated from C57BL/6xC3H E14 embryonic mouse livers, as described in (Strick-Marchand and Weiss, 2002). Briefly, livers at 14 dpc were separately dissected, and cell suspension was plated onto collagen-coated 100-mm petri dishes in hepatocyte attachment medium (Invitrogen, Cergy Pontoise Cedex, France). The next day and thereafter, medium was replaced by RPMI 1640 (Eurobio) containing 10% fetal bovine serum (Eurobio), 50 ng/ml epidermal growth factor, 30 ng/ml insulin-like growthfactor II (PeproTech, Rocky Hill, NJ), 10 mg/ml insulin (Sigma) and antibiotics. Clones were picked after 2–3 months of culture and expanded in the same medium on dishes coated with Collagen I (BD Biosciences, Le Pont de Claix, France) in humidified atmosphere with 5% CO2 at 37°C.

### Isolation of cell populations

Human H-Ras^G12V^ cDNA sequence was cloned into LeGO-iV2 and LeGO-iC2 bicistronic vectors (Addgene #27344 and #27345, respectively). Retroviruses were produced by JetPEI (Polypus-Transfection) transfection of HEK293T cells with HRas^G12V^-iV2 and HRas^G12V^-iC2 constructs together with Gag/pol and pCAG-Eco packaging vectors. Culture media containing viruses were filtered (0,45 μm) and used to infect BMEL cells. Based on Venus (530/30nm) and mCherry (610/20nm) fluorescence intensity, Aria IIU and IIIU cell sorters (Becton Dickinson) were used to isolate HRas^G12V^-LOW and HRas^G12V^-HIGH populations from HRas^G12V^-iV2 and HRas^G12V^-iC2 BMEL cells.

### Soft agar assay

10^5^ cells/well were seeded in 0.5% agar diluted in RPMI media. 1 mL media was added to the agar layer and was changed every 3 days. After 3 weeks, colonies were stained with 0,005% crystal violet 4% PFA. Colonies were counted automatically in imageJ using the “Analyze particles” function.

### Orthotopic xenografts

Athymic nude (Crl:NU(NCr)-Foxn1nu), NOD SCID gamma (NOD.Cg-PrkdcSCID Il2rgtm1Wjl/SzJ) or C57BL/6J mice from Charles River were anesthetized using intraperitoneal injections of Ketamine/Xylazine. A 1,5cm transversal incision of the skin was performed bellow the sternum’s xyphoid process and followed by a transversal incision of the peritoneum, the tip of the liver left lateral lobe was pulled out of the peritoneal cavity. BMEL cells suspension in PBS – 20% Matrigel (Corning) were injected in the liver parenchyma. Stitching of the peritoneum then skin was performed using surgical suture (Monosof 5/0 3/8C 16mm).

### Analysis of tumour cellular composition

Tumors were fixed in 4% PFA for 4 hours and left overnight in 30% sucrose solution. Fixed tissues were placed in OCT cryogenic matrix and frozen in liquid nitrogen. Sections were cut at −20°C and mounted on superFROST slides with ProlonGold mounting media. Whole slide scans were performed with Zeiss Axioscan at 20X magnification and DAPI (405nm), GFP (488nm) and Cy3 (550nm) fluorescent channels. On average 10 sections from different regions of the tumours were analysed per mouse. Quantification was done by delimiting GFP positive or Cy3 positive regions and measuring their area on ZEN software. Alternatively, tumours were dissociated with mouse tumour dissociation kit (Miltenyi) and analysed with Novocyte ACEA flow cytometer using FITC (530/30BP) and mCherry (615/20BP) filter cubes.

### RNA extraction and quantification

Total RNA was isolated from cells, dissociated and sorted tumors or frozen tissue samples using RNeasy mini kit (Qiagen). cDNA was prepared from 500ng of total RNA using QuantiTect Reverse Transcription Kit (Qiagen). SYBR green (see primers list) or Taqman based Quantitative PCR (hprt: Roche universal probe library #95 (agtcccag); Ras: Roche universal probe library #88 (catcctcc); Ceacam1: (tctcacagagcacaaaccctcagc) were performed on Roche LightCycler480.

### Primers list

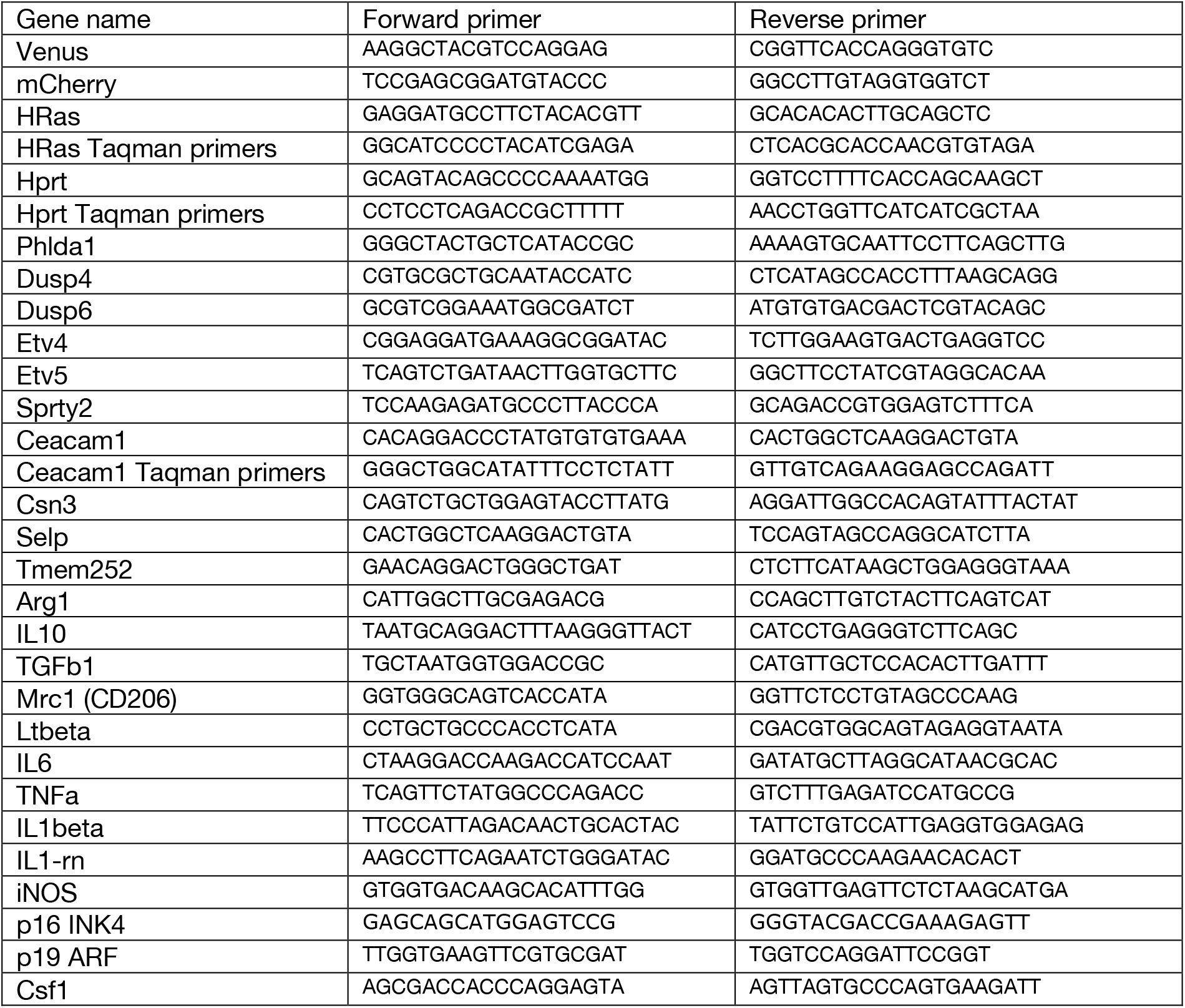

### RNAseq analysis

RNAseq bank were prepared from, respectively, 3 and 4 independent biological samples for the cell lines and Venus-positive cells sorted from tumours. We used *Universal Plus mRNAseq* kit from NuGEN and sequenced on Illumina HiSeq 2500. After quality control the sequences were aligned to Mm9 mouse genome using TopHat2. Normalization and differential gene expression between pairs was performed with Deseq. Gene set enrichment analysis was done using pre-ranked method based on fold changed values (GSEAv.6.0) and analysed with C5 ontology gene sets (Subramanian et al., 2005). Immune cell type deconvolution was done with CIBERSORTx (https://cibersortx.stanford.edu/) based on RPKM counts that were compared with murine immune signatures from 10 cell types produced by (Chen et al., 2017).

### Flow Cytometry analyses

Mouse tumors were dissociated into single-cell suspension using the Mouse tumour dissociation kit (Miltenyi Biotec) and the gentleMACS^™^ Octo Dissociator following manufacturer’s instructions and as described in (Taranto et al., 2021). The cell suspension was passed through a 70*μ*m cell strainer (Miltenyi), centrifuged at 300g for 10 min at 4°C and washed 3 times in FACS buffer. GFP negative stromal cells were isolated from cell suspensions with a FACS sorter ARIAIIu^®^ (Becton Dickinson) and frozen in foetal calf serum with 10% DMSO. Frozen samples were thawed under sterile conditions and single cell preparations were incubated with anti-CD16/CD32 antibody (BD Biosciences) for 15 minutes and stained with the indicated antibodies following standard procedures. Samples were fixed with eBioscience fixation/permeabilization kit (Invitrogen) and Ki67 antibody was used for intracellular staining (*see Table below)*, and analysed as previously described (Wang et al., 2019). The signal was detected by a 4 laser Fortessa^®^ flow cytometer (Becton Dickinson). Analyses were carried out using FlowJo^®^ software.

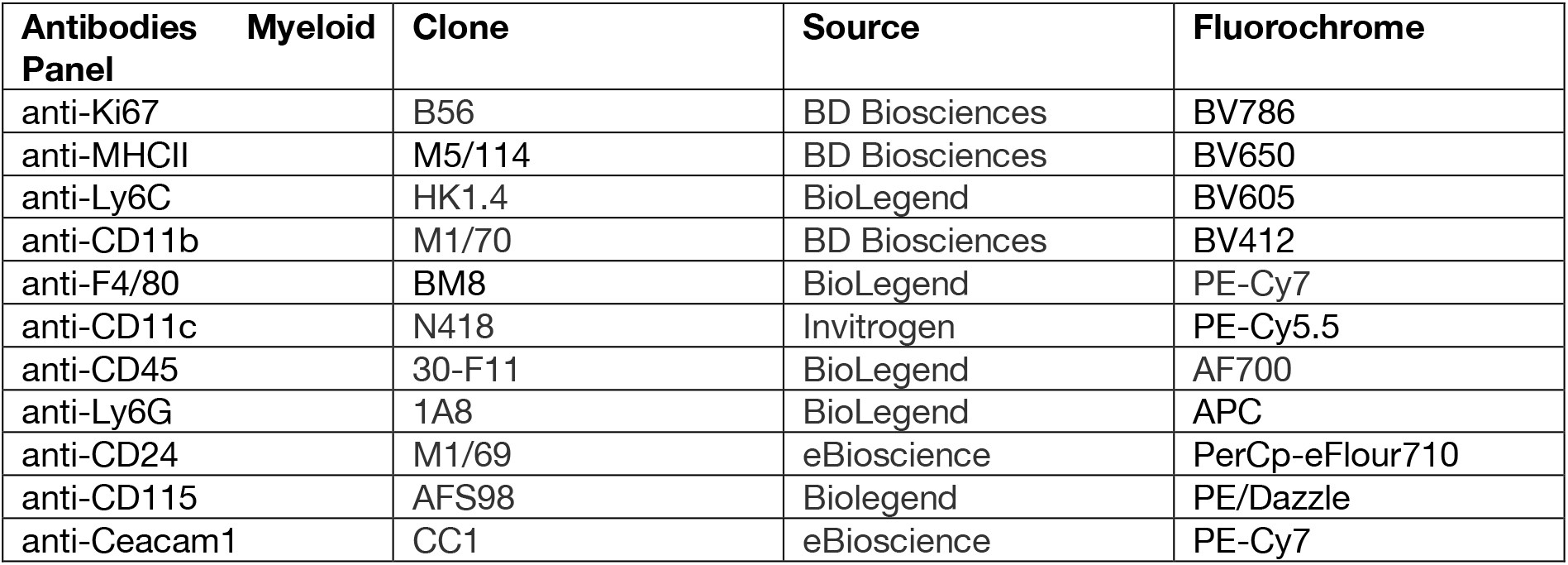

### Immunohistochemical analysis

Antibodies used:

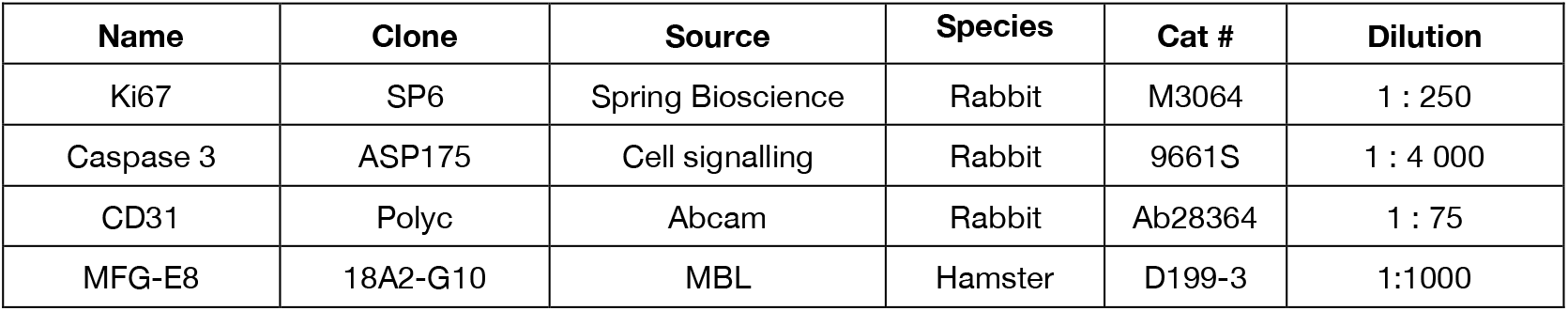

Immunohistochemical analysis Antibodies used:
| Name | Clone | Source | Species | Cat # | Dilution |
| --- | --- | --- | --- | --- | --- |
| Ki67 | SP6 | Spring Bioscience | Rabbit | M3064 | 1 : 250 |
| Caspase 3 | ASP175 | Cell signalling | Rabbit | 9661S | 1 : 4 000 |
| CD31 | Polyc | Abcam | Rabbit | Ab28364 | 1 : 75 |
| MFG-E8 | 18A2-G10 | MBL | Hamster | D199-3 | 1:1000 |

Three μm sections of paraffin-embedded samples were analyzed using a Ventana Discovery Ultra automated staining instrument (Ventana Medical Systems), according to the manufacturer’s instructions. Briefly, slides were de-paraffinized, the epitope retrieval was performed with the reagents provided by the manufacturer, the endogenous peroxidase was blocked and the samples were incubated with the appropriate antibodies for 60 min at 37°C. Signal enhancement for Caspase 3 and Ki67 antibodies was performed either with the OmniMap anti-rabbit detection kit or by the Discovery HQ conjugated anti-rabbit IgG followed by Discovery amplification anti-HQ HRP Multimer, according to the manufacturer’s instructions.

Slides were incubated with DAB chromogen and counterstained with hematoxylin II for 8 min, followed by Bluing reagent for 4 min. Brightfield stained slides were digitalized with a Hamamatsu NanoZoomer 2.0-HT scanner and images were visualized with the NDP view 1.2.47 software except for the MFG-E8 staining, for which slides were digitally processed using the Aperio ScanScope (Aperio). Nodule size was drawn by hand in HALO^™^ image analysis software (Indica Labs) and an algorithm was designed with the Multiplex IHC v1.2 module to quantify the percentage of positive cells per mouse, as indicated in figure legends.

### Immunofluorescence

Frozen sections were equilibrated at room temperature for 10min and rehydrated in PBS. Sections were incubated overnight with anti-mouse NKp46/NRC1 antibody (1:500, AF2225 R&D system) and for 1h with rabbit anti-goat alexa647 conjugated secondary antibody (Invitrogen A21446). Images were acquired using Zeiss axioimager Z1 microscope with DAPI, GFP and Cy5 filter cubes

### Software

The following open-source code and software were used in this study: R (v.3.5.1), Default settings were used for all GSEA analyses. R package DESeq2 (v.1.22.1), R package heatmap (v.1.0.10) and R package VennDiagram (v.1.6.20) were also used. Graphpad prism v8 was used to make graphs and for statistical analyses.

## Data availability

The RNA-sequencing data have been deposited in the Gene Expression Omnibus (GEO, NCBI) repository, and are accessible through GEO Series accession number GSE180580.

## Acknowledgments

This work was supported by the grant HTE from ITMO Cancer (to UH), from the Association Française pour l’Etude du Foie (AFEF, to DG) and from SIRIC Montpellier Cancer (to UH). AL received support from the University of Montpellier and from the Fondation ARC and SIRIC Montpellier Cancer. LA, SV and CR are supported by the Dutch Cancer Society (KWF 12049/2018-2), Oncode Institute and the Center for Cancer Genomics (CGC.nl). We acknowledge the “Réseau d’Histologie Expérimentale de Montpellier” - RHEM facility, supported by SIRIC Montpellier Cancer (Grant INCa_Inserm_DGOS_12553), the European regional development foundation and the Occitanie region (FEDER-FSE 2014-2020 Languedoc Roussillon), for the histological analyses. We acknowledge the MGX facility for the RNAseq analysis, and the MRI imaging platform. We thank the RAM animal facility and IGMM zootechnicians for care of animals. We are especially grateful to Myriam Boyer and Stéphanie Viala for help with cell sorting. We thank José Ursic-Bedoya for the cell line derived from the hydrodynamic gene transfer experiments and Benjamin Rivière for his expert advice on anatomopathology. We gratefully acknowledge many helpful discussions with Eric Assenat and all other members of the Hibner laboratory. We are especially grateful to the direction of the IGMM institute for ensuring the conditions that allowed us to continue our work in the past year.

## Authors contributions

AL: Conceptualization, Investigation, Visualization, Writing-review and editing; FRS: Investigation; CR: Investigation; SV: Investigation; GD: Investigation, Formal analysis, Visualization; AR: Investigation; VD: Investigation; AZEA: Formal analysis; PF: Formal analysis, Writing-review and editing; LA: Supervision, Funding acquisition, Writing-review and editing; UH: Conceptualization, Supervision, Funding acquisition, Writing-original draft, Writing-review and editing. DG: Conceptualization, Supervision, Investigation, Funding acquisition, Visualization, Writing-original draft, Writing-review and editing.

## Bibliography

Akkari, L., Haouzi, D., Binamé, F., Floc’h, N., Lassus, P., Baghdiguian, S., and Hibner, U. (2010). Cell shape and TGF-β signaling define the choice of lineage during in vitro differentiation of mouse primary hepatic precursors. J. Cell. Physiol. 225, 186–195.

Akkari, L., Grégoire, D., Floc’h, N., Moreau, M., Hernandez, C., Simonin, Y., Rosenberg, A.R., Lassus, P., and Hibner, U. (2012). Hepatitis C viral protein NS5A induces EMT and participates in oncogenic transformation of primary hepatocyte precursors. J. Hepatol. 57, 1021–1028.

Ambrogio, C., Köhler, J., Zhou, Z.-W., Wang, H., Paranal, R., Li, J., Capelletti, M., Caffarra, C., Li, S., Lv, Q., et al. (2018). KRAS Dimerization Impacts MEK Inhibitor Sensitivity and Oncogenic Activity of Mutant KRAS. Cell 172, 857–868.e15.

Bacevic, K., Prieto, S., Caruso, S., Camasses, A., Dubra, G., Ursic-Bedoya, J., Lozano, A., Butterworth, J., Zucman-Rossi, J., Hibner, U., et al. (2019). CDK8 and CDK19 kinases have non-redundant oncogenic functions in hepatocellular carcinoma. BioRxiv 789586.

Baghdadi, M., Chiba, S., Yamashina, T., Yoshiyama, H., and Jinushi, M. (2012). MFG-E8 Regulates the Immunogenic Potential of Dendritic Cells Primed with Necrotic Cell-Mediated Inflammatory Signals. PLOS ONE 7, e39607.

Brant, R., Sharpe, A., Liptrot, T., Dry, J.R., Harrington, E.A., Barrett, J.C., Whalley, N., Womack, C., Smith, P., and Hodgson, D.R. (2017). Clinically Viable Gene Expression Assays with Potential for Predicting Benefit from MEK Inhibitors. Clin. Cancer Res. Off. J. Am. Assoc. Cancer Res. 23, 1471–1480.

Bremner, R., and Balmain, A. (1990). Genetic changes in skin tumor progression: Correlation between presence of a mutant ras gene and loss of heterozygosity on mouse chromosome 7. Cell 61, 407–417.

Chan, L.-K., Ho, D.W.-H., Kam, C.S., Chiu, E.Y.-T., Lo, I.L.-O., Yau, D.T.-W., Cheung, E.T.-Y., Tang, C.-N., Tang, V.W.-L., Lee, T.K.-W., et al. (2021). RSK2-inactivating mutations potentiate MAPK signaling and support cholesterol metabolism in hepatocellular carcinoma. J. Hepatol. 74, 360–371.

Chen, Z., Huang, A., Sun, J., Jiang, T., Qin, F.X.-F., and Wu, A. (2017). Inference of immune cell composition on the expression profiles of mouse tissue. Sci. Rep. 7, 40508.

Dankner, M., Gray-Owen, S.D., Huang, Y.-H., Blumberg, R.S., and Beauchemin, N. (2017). CEACAM1 as a multi-purpose target for cancer immunotherapy. Oncoimmunology 6.

Delire, B., and Stärkel, P. (2015). The Ras/MAPK pathway and hepatocarcinoma: pathogenesis and therapeutic implications. Eur. J. Clin. Invest. 45, 609–623.

Dikic, I., Schlessinger, J., and Lax, I. (1994). PC12 cells overexpressing the insulin receptor undergo insulin-dependent neuronal differentiation. Curr. Biol. 4, 702–708.

Dorard, C., Vucak, G., and Baccarini, M. (2017). Deciphering the RAS/ERK pathway in vivo. Biochem. Soc. Trans. 45, 27–36.

Dou, L., Shi, X., He, X., and Gao, Y. (2020). Macrophage Phenotype and Function in Liver Disorder. Front. Immunol. 10.

East, P., Kelly, G.P., Biswas, D., Marani, M., Hancock, D.C., Creasy, T., Sachsenmeier, K., Swanton, C., Consortium, on behalf of the Tracer., Trécesson, S. de C., et al. (2021). Oncogenic RAS activity predicts response to chemotherapy and outcome in lung adenocarcinoma. BioRxiv 2021.04.02.437896.

Gerstung, M., Jolly, C., Leshchiner, I., Dentro, S.C., Gonzalez, S., Rosebrock, D., Mitchell, T.J., Rubanova, Y., Anur, P., Yu, K., et al. (2020). The evolutionary history of 2,658 cancers. Nature 578, 122–128.

Gonzaga, R., Matzinger, P., and Perez-Diez, A. (2011). Resident Peritoneal NK Cells. J. Immunol. 187, 6235–6242.

Gross, S.M., Dane, M.A., Bucher, E., and Heiser, L.M. (2019). Individual Cells Can Resolve Variations in Stimulus Intensity along the IGF-PI3K-AKT Signaling Axis. Cell Syst. 9, 580–588.e4.

Hallberg, B., Rayter, S.I., and Downward, J. (1994). Interaction of Ras and Raf in intact mammalian cells upon extracellular stimulation. J. Biol. Chem. 269, 3913–3916.

Helfrich, I., and Singer, B.B. (2019). Size Matters: The Functional Role of the CEACAM1 Isoform Signature and Its Impact for NK Cell-Mediated Killing in Melanoma. Cancers 11.

Ischenko, I., D’Amico, S., Rao, M., Li, J., Hayman, M.J., Powers, S., Petrenko, O., and Reich, N.C. (2021). KRAS drives immune evasion in a genetic model of pancreatic cancer. Nat. Commun. 12, 1482.

Janiszewska, M., Tabassum, D.P., Castaño, Z., Cristea, S., Yamamoto, K.N., Kingston, N.L., Murphy, K.C., Shu, S., Harper, N.W., Del Alcazar, C.G., et al. (2019). Subclonal cooperation drives metastasis by modulating local and systemic immune microenvironments. Nat. Cell Biol. 21, 879–888.

Katyal, S., Oliver, J.H., Peterson, M.S., Ferris, J.V., Carr, B.S., and Baron, R.L. (2000). Extrahepatic metastases of hepatocellular carcinoma. Radiology 216, 698–703.

Kerr, E.M., Gaude, E., Turrell, F.K., Frezza, C., and Martins, C.P. (2016). Mutant Kras copy number defines metabolic reprogramming and therapeutic susceptibilities. Nature 531, 110–113.

Kortlever, R.M., Sodir, N.M., Wilson, C.H., Burkhart, D.L., Pellegrinet, L., Brown Swigart, L., Littlewood, T.D., and Evan, G.I. (2017). Myc Cooperates with Ras by Programming Inflammation and Immune Suppression. Cell 171, 1301–1315.e14.

Kumagai, Y., Naoki, H., Nakasyo, E., Kamioka, Y., Kiyokawa, E., and Matsuda, M. (2015). Heterogeneity in ERK activity as visualized by in vivo FRET imaging of mammary tumor cells developed in MMTV-Neu mice. Oncogene 34, 1051–1057.

Lenormand, P., Sardet, C., Pagès, G., L’Allemain, G., Brunet, A., and Pouysségur, J. (1993). Growth factors induce nuclear translocation of MAP kinases (p42mapk and p44mapk) but not of their activator MAP kinase kinase (p45mapkk) in fibroblasts. J. Cell Biol. 122, 1079–1088.

Lim, H.Y., Merle, P., Weiss, K.H., Yau, T., Ross, P., Mazzaferro, V., Blanc, J.-F., Ma, Y.T., Yen, C.J., Kocsis, J., et al. (2018). Phase II Studies with Refametinib or Refametinib plus Sorafenib in Patients with RAS-Mutated Hepatocellular Carcinoma. Clin. Cancer Res. Off. J. Am. Assoc. Cancer Res. 24, 4650–4661.

Llovet, J.M., Kelley, R.K., Villanueva, A., Singal, A.G., Pikarsky, E., Roayaie, S., Lencioni, R., Koike, K., Zucman-Rossi, J., and Finn, R.S. (2021). Hepatocellular carcinoma. Nat. Rev. Dis. Primer 7, 1–28.

Losic, B., Craig, A.J., Villacorta-Martin, C., Martins-Filho, S.N., Akers, N., Chen, X., Ahsen, M.E., von Felden, J., Labgaa, I., D’Avola, D., et al. (2020). Intratumoral heterogeneity and clonal evolution in liver cancer. Nat. Commun. 11, 291.

Marshall, C.J. (1995). Specificity of receptor tyrosine kinase signaling: transient versus sustained extracellular signal-regulated kinase activation. Cell 80, 179–185.

Marusyk, A., Janiszewska, M., and Polyak, K. (2020). Intratumor Heterogeneity: The Rosetta Stone of Therapy Resistance. Cancer Cell 37, 471–484.

Molina-Sánchez, P., Ruiz de Galarreta, M., Yao, M.A., Lindblad, K.E., Bresnahan, E., Bitterman, E., Martin, T.C., Rubenstein, T., Nie, K., Golas, J., et al. (2020). Cooperation Between Distinct Cancer Driver Genes Underlies Intertumor Heterogeneity in Hepatocellular Carcinoma. Gastroenterology 159, 2203–2220.e14.

Mueller, S., Engleitner, T., Maresch, R., Zukowska, M., Lange, S., Kaltenbacher, T., Konukiewitz, B., Öllinger, R., Zwiebel, M., Strong, A., et al. (2018). Evolutionary routes and KRAS dosage define pancreatic cancer phenotypes. Nature 554, 62–68.

Murphy, L.O., and Blenis, J. (2006). MAPK signal specificity: the right place at the right time. Trends Biochem. Sci. 31, 268–275.

Newman, A.M., Steen, C.B., Liu, C.L., Gentles, A.J., Chaudhuri, A.A., Scherer, F., Khodadoust, M.S., Esfahani, M.S., Luca, B.A., Steiner, D., et al. (2019). Determining cell type abundance and expression from bulk tissues with digital cytometry. Nat. Biotechnol. 37, 773–782.

Pagès, G., Lenormand, P., L’Allemain, G., Chambard, J.C., Meloche, S., and Pouysségur, J. (1993). Mitogen-activated protein kinases p42mapk and p44mapk are required for fibroblast proliferation. Proc. Natl. Acad. Sci. U. S. A. 90, 8319–8323.

Pastushenko, I., and Blanpain, C. (2019). EMT Transition States during Tumor Progression and Metastasis. Trends Cell Biol. 29, 212–226.

Peng, Y., and Elkon, K.B. (2012). Autoimmunity in MFG-E8-deficient mice is associated with altered trafficking and enhanced crosspresentation of apoptotic cell antigens. J. Clin. Invest. 122, 782–782.

Pouysségur, J., Volmat, V., and Lenormand, P. (2002). Fidelity and spatio-temporal control in MAP kinase (ERKs) signalling. Biochem. Pharmacol. 64, 755–763.

Ringelhan, M., Pfister, D., O’Connor, T., Pikarsky, E., and Heikenwalder, M. (2018). The immunology of hepatocellular carcinoma. Nat. Immunol. 19, 222–232.

Sällberg, M., and Pasetto, A. (2020). Liver, Tumor and Viral Hepatitis: Key Players in the Complex Balance Between Tolerance and Immune Activation. Front. Immunol. 11.

Sanchez-Vega, F., Mina, M., Armenia, J., Chatila, W.K., Luna, A., La, K.C., Dimitriadoy, S., Liu, D.L., Kantheti, H.S., Saghafinia, S., et al. (2018). Oncogenic Signaling Pathways in The Cancer Genome Atlas. Cell 173, 321–337.e10.

Serrano, M., Lin, A.W., McCurrach, M.E., Beach, D., and Lowe, S.W. (1997). Oncogenic ras provokes premature cell senescence associated with accumulation of p53 and p16INK4a. Cell 88, 593–602.

Shi, F.-D., Ljunggren, H.-G., La Cava, A., and Van Kaer, L. (2011). Organ-specific features of natural killer cells. Nat. Rev. Immunol. 11, 658–671.

Sodir, N.M., Kortlever, R.M., Barthet, V.J.A., Campos, T., Pellegrinet, L., Kupczak, S., Anastasiou, P., Swigart, L.B., Soucek, L., Arends, M.J., et al. (2020). MYC Instructs and Maintains Pancreatic Adenocarcinoma Phenotype. Cancer Discov. 10, 588–607.

Strick-Marchand, H., and Weiss, M.C. (2002). Inducible differentiation and morphogenesis of bipotential liver cell lines from wild-type mouse embryos. Hepatol. Baltim. Md 36, 794–804.

Strick-Marchand, H., and Weiss, M.C. (2003). Embryonic liver cells and permanent lines as models for hepatocyte and bile duct cell differentiation. Mech. Dev. 120, 89–98.

Strick-Marchand, H., Morosan, S., Charneau, P., Kremsdorf, D., and Weiss, M.C. (2004). Bipotential mouse embryonic liver stem cell lines contribute to liver regeneration and differentiate as bile ducts and hepatocytes. Proc. Natl. Acad. Sci. U. S. A. 101, 8360–8365.

Subramanian, A., Tamayo, P., Mootha, V.K., Mukherjee, S., Ebert, B.L., Gillette, M.A., Paulovich, A., Pomeroy, S.L., Golub, T.R., Lander, E.S., et al. (2005). Gene set enrichment analysis: A knowledge-based approach for interpreting genome-wide expression profiles. Proc. Natl. Acad. Sci. 102, 15545–15550.

Suda, T., Tatsumi, T., Nishio, A., Kegasawa, T., Yoshioka, T., Yamada, R., Furuta, K., Kodama, T., Shigekawa, M., Hikita, H., et al. (2018). CEACAM1 Is Associated With the Suppression of Natural Killer Cell Function in Patients With Chronic Hepatitis C. Hepatol. Commun. 2, 1247–1258.

Taranto, D., Ramirez, C.F., Vegna, S., de Groot, M.H., de Wit, N., Van Baalen, M., Klarenbeek, S., and Akkari, L. (2021). Multiparametric analyses of hepatocellular carcinoma somatic mouse models and their associated tumor microenvironment. Curr. Protoc. e147.

Traverse, S., Seedorf, K., Paterson, H., Marshall, C.J., Cohen, P., and Ullrich, A. (1994). EGF triggers neuronal differentiation of PC12 cells that overexpress the EGF receptor. Curr. Biol. 4, 694–701.

Wellenstein, M.D., and de Visser, K.E. (2018). Cancer-Cell-Intrinsic Mechanisms Shaping the Tumor Immune Landscape. Immunity 48, 399–416.

Woods, D., Parry, D., Cherwinski, H., Bosch, E., Lees, E., and McMahon, M. (1997). Raf-induced proliferation or cell cycle arrest is determined by the level of Raf activity with arrest mediated by p21Cip1. Mol. Cell. Biol. 17, 5598–5611.

Xue, W., Zender, L., Miething, C., Dickins, R.A., Hernando, E., Krizhanovsky, V., Cordon-Cardo, C., and Lowe, S.W. (2007). Senescence and tumour clearance is triggered by p53 restoration in murine liver carcinomas. Nature 445, 656–660.

Yu, W.-J., Yang, W.-H., Shi, Z.-X., Yang, X.-D., and Wang, H.-J. (2008). [Application of NOD/SCID mice in research of experimental hematology - review]. Zhongguo Shi Yan Xue Ye Xue Za Zhi 16, 964–968.

Zucman-Rossi, J., Villanueva, A., Nault, J.-C., and Llovet, J.M. (2015). Genetic Landscape and Biomarkers of Hepatocellular Carcinoma. Gastroenterology 149, 1226–1239.e4.

Zugasti, O., Rul, W., Roux, P., Peyssonnaux, C., Eychene, A., Franke, T.F., Fort, P., and Hibner, U. (2001). Raf-MEK-Erk cascade in anoikis is controlled by Rac1 and Cdc42 via Akt. Mol. Cell. Biol. 21, 6706–6717.

