## Supplemental data for "Ras/MAPK signalling intensity defines subclonal fitness in a mouse model of primary and metastatic hepatocellular carcinoma"

### Figure S1

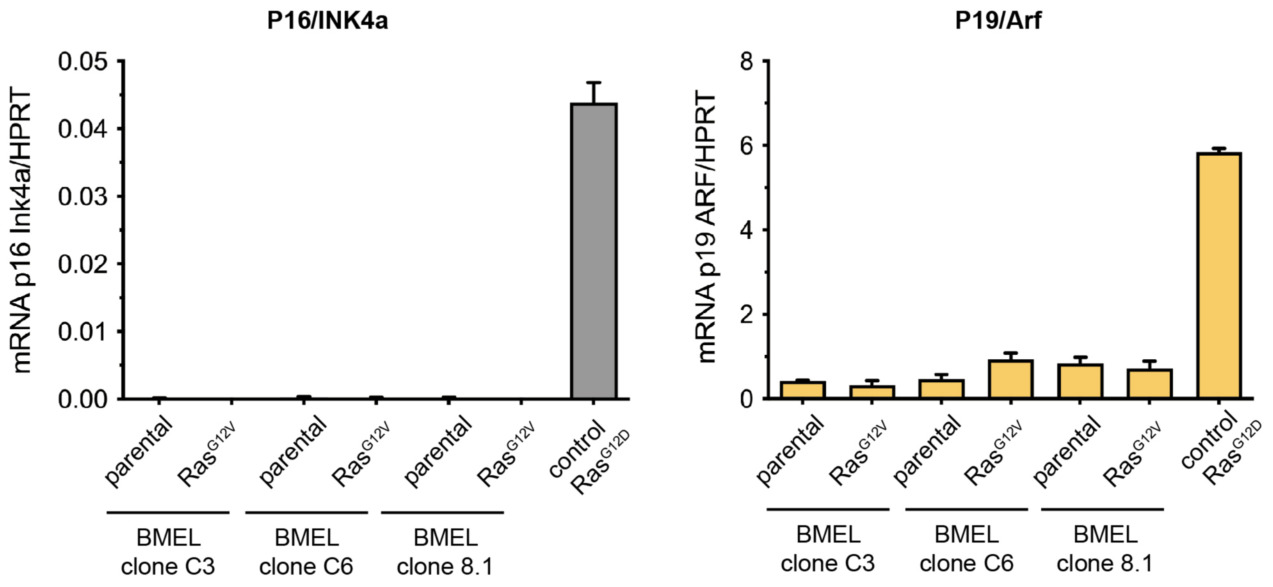

**Suppl. Fig. 1: BMEL clones do not express the p16INK4A tumour suppressor.** Quantification of mRNA expression levels of p16INK4A and p19ARF using qRT-PCR performed on three independent BMEL clones (C3, C6 and 8.1) with or without oncogenic Ras expression, as indicated. $NRAS^{G12D}$ p53null tumour cell line derived from a mouse tumour following a hydrodynamic gene transfer is used as control (control $Ras^{G12D}$).

### Figure S2

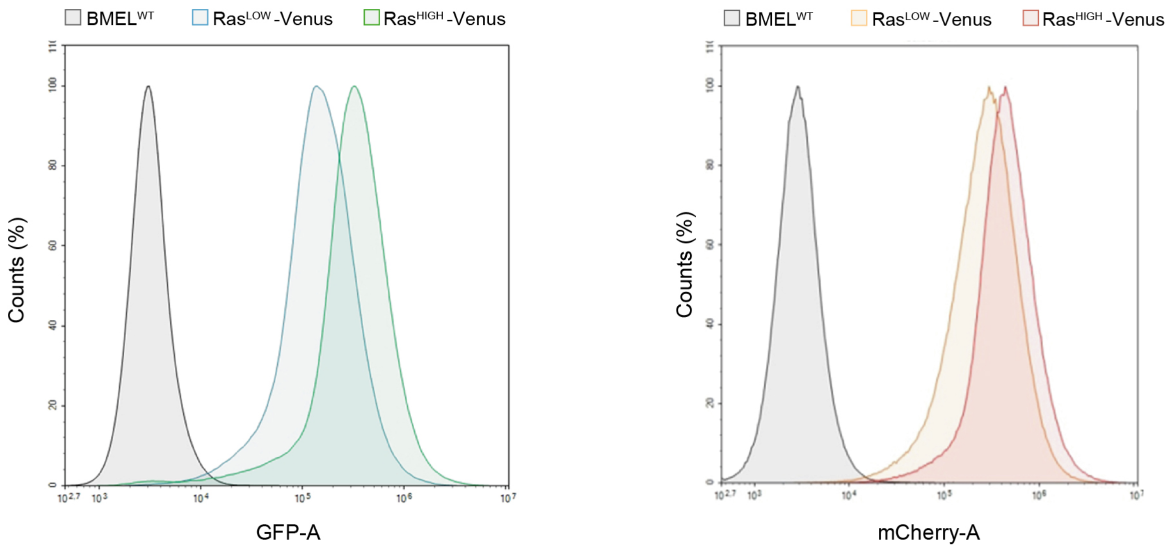

**Suppl. Fig. 2: Overlap in transgene expression levels in sorted cell populations.** Fluorescence intensity profiles of «$Ras^{LOW}$» and «$Ras^{HIGH}$» Venus (left panel) and mCherry (right panel) sorted BMEL cell populations.

### Figure S3

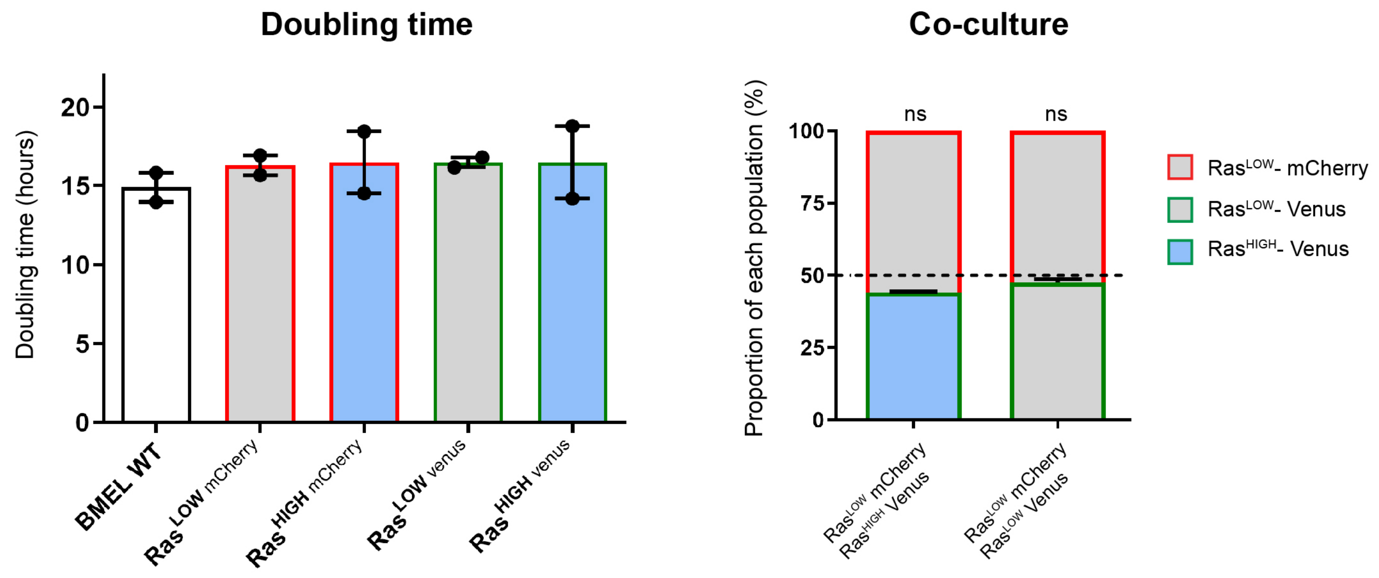

**Suppl Fig 3 - Ras$^{G12V}$ expression level does not change BMEL proliferation rates *ex vivo*.** Left panel shows doubling times in 2D culture for the indicated cell lines. Mean and values from two independent experiments are shown. Right panel shows the composition of cellular populations after 5 days of culture of 1:1 mix of Ras$^{HIGH}$-Venus and Ras$^{LOW}$-mCherry cells and 1:1 mix of Ras$^{LOW}$-Venus and Ras$^{LOW}$-mCherry cells. Mean +/- SEM are indicated for three independent experiments. Deviations from the theoretical value of 50% (dashed line) were calculated using one sample t-test.

#### Figure S4

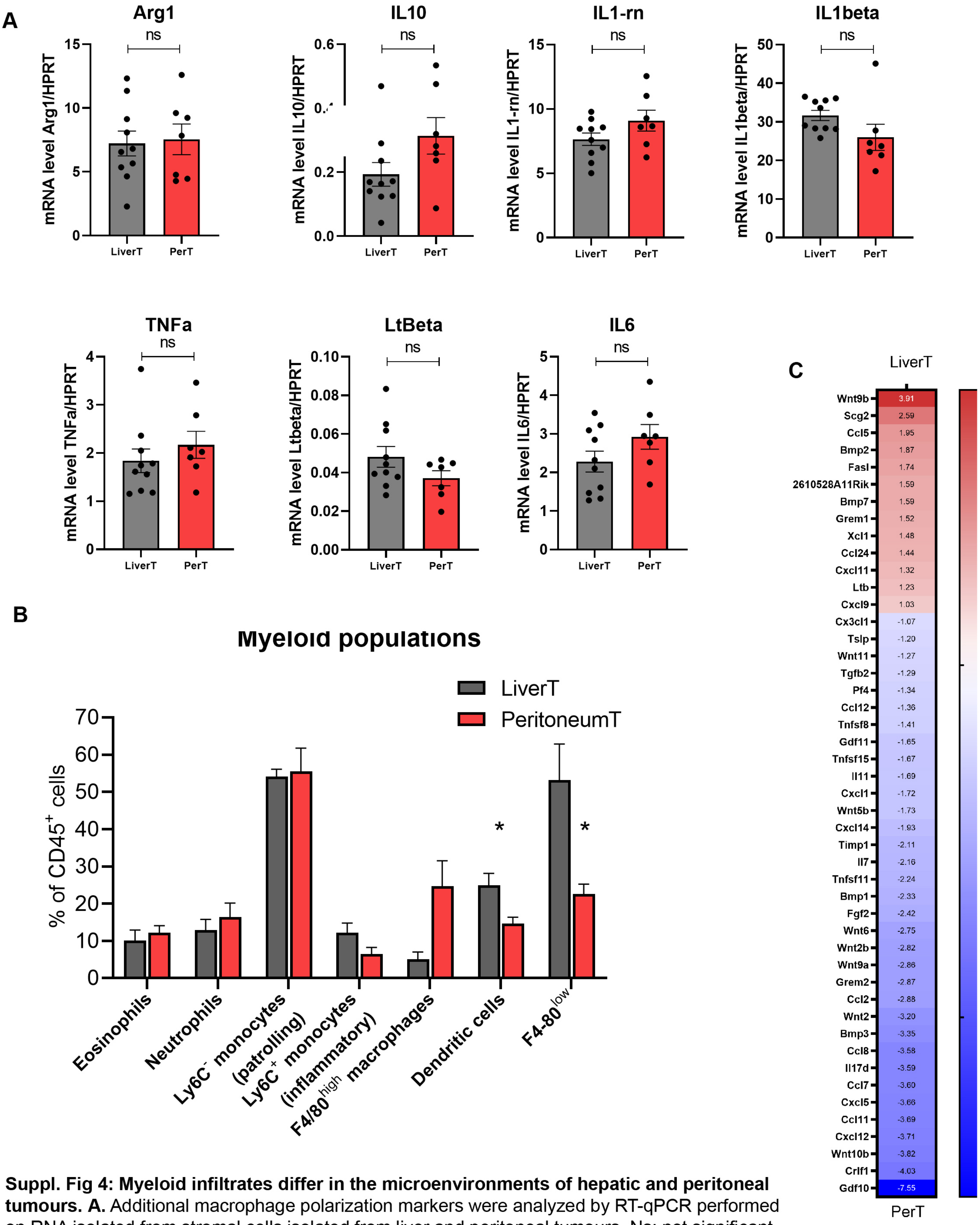

**Suppl. Fig 4: Myeloid infiltrates differ in the microenvironments of hepatic and peritoneal tumours. A.** Additional macrophage polarization markers were analyzed by RT-qPCR performed on RNA isolated from stromal cells isolated from liver and peritoneal tumours. Ns: not significant **B.** Quantification of myeloid populations defined as $CD45^{+}CD11b^{+}Ly6G^{int}Ly6C^{int}$ eosinophils, $CD45^{+}CD11b^{+}Ly6G^{+}$ neutrophils, $CD45^{+}CD11b^{+}Ly6G^{-}Ly6C^{-}$ patrolling monocytes, $CD45^{+}CD11b^{+}Ly6G^{-}Ly6C^{+}$inflammatory monocytes, $CD45^{+}CD11b^{+}Ly6G^{-}Ly6C^{-}F4/80^{high}$ macrophages, $CD45^{+}CD11c^{+}MHCII^{high}F4/80^{-}$ $CD24^{+}$ dendritic cells and $F4/80^{low}$ immature macrophages by flow cytometry analysis of sorted liver and peritoneal stromal cells from athymic nude mice tumours (n=4). Student t-test, p-value: *<0.05 . **C.** Heatmap of cytokines differentially expressed in stromal cells of liver and peritoneal tumours. Numbers display Log2 fold change values for each gene (Deseq, Log2FC>1.5 pvalue<0.05).

**Figure S5**

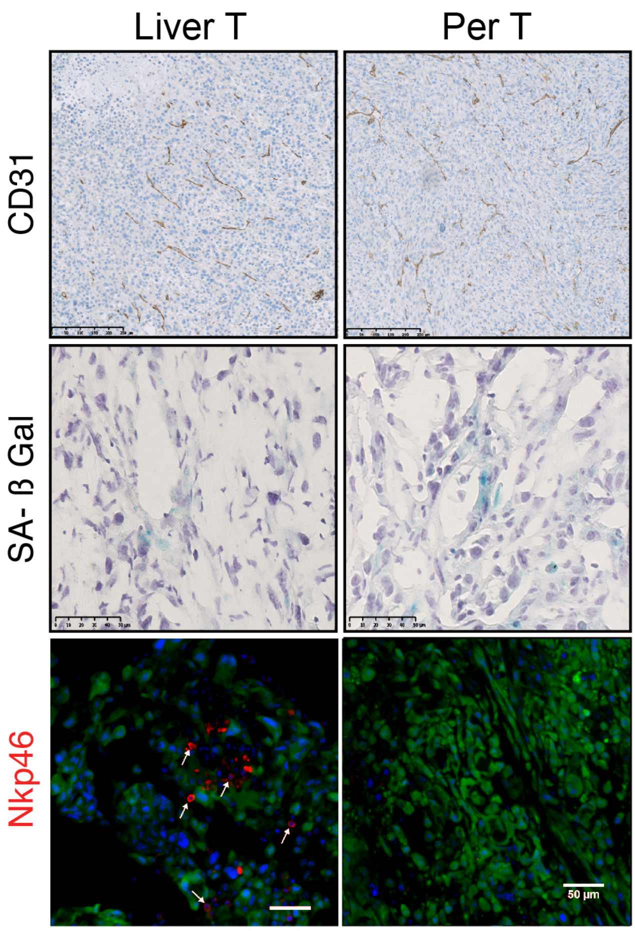

**Suppl. Fig. 5 : Differences in NK cells infiltration between liver and peritoneal tumours.** Immunhistochemistry performed on athymic nude mice liver and peritoneal tumours initiated by inoculation of Ras$^{HIGH}$ BMEL cells were analyzed in respect to anti-CD31 (upper panels) and senescence-associated beta-galatosidase staining (middle panels). Lower panels: immuno-fluorescence staining of Ras$^{HIGH}$ (Venus-labelled, green) tumours with the NK-specific anti-Nkp46 antibody (red). Arrows show Nkp46 positive cells that infiltrated liver tumours.

**Supplementary Table 1: gene signature for the low threshold Ras/MAPK signalling (Figure 1E left panel)**

| n=160 | | WT vs $Ras^{LOW}$ | | WT vs $Ras^{HIGH}$ | | $Ras^{LOW}$ vs $Ras^{HIGH}$ | |
|---|---|---|---|---|---|---|---|
| Gene symbol | Geneid | log2FoldChange | padjExactTest | log2FoldChange | padjExactTest | log2FoldChange | padjExactTest |
| Lgals2 | *107753* | 1,51 | 4,22E-08 | 1,66 | 8,78E-10 | 0,67 | 1,00E+00 |
| Rap1b | *215449* | 3,17 | 2,44E-14 | 3,97 | 1,61E-46 | 0,01 | 3,92E-01 |
| Nup107 | *103468* | 3,59 | 3,38E-05 | 3,99 | 6,45E-06 | 0,47 | 1,00E+00 |
| Hmga1 | *15361* | 2,13 | 2,30E-07 | 2,64 | 2,63E-03 | 0,26 | 1,00E+00 |
| Rbp4 | *19662* | 2,03 | 3,40E-14 | 2,50 | 3,31E-05 | 0,15 | 1,00E+00 |
| Slc35e3 | *215436* | 3,02 | 3,70E-29 | 3,57 | 1,62E-38 | 0,09 | 1,00E+00 |
| Ankrd1 | *107765* | -1,65 | 3,59E-04 | -2,70 | 2,09E-07 | 0,01 | 3,51E-01 |
| Ccn1 | *16007* | -1,86 | 3,56E-03 | -3,06 | 2,15E-06 | 0,08 | 9,78E-01 |
| Sprr1a | *20753* | -2,23 | 5,75E-04 | -1,62 | 4,69E-02 | 0,47 | 1,00E+00 |
| Prkg2 | *19092* | 1,82 | 7,48E-05 | 2,36 | 1,85E-18 | 0,11 | 1,00E+00 |
| Phlda2 | *22113* | 2,57 | 9,07E-04 | 3,11 | 1,58E-04 | 0,32 | 1,00E+00 |
| Peg3os | *100169889* | -2,44 | 6,38E-03 | -1,71 | 2,04E-02 | 0,41 | 1,00E+00 |
| Psph | *100678* | 1,71 | 1,75E-03 | 2,19 | 3,16E-14 | 0,15 | 1,00E+00 |
| Rps6ka2 | *20112* | 2,30 | 1,64E-16 | 3,33 | 7,56E-06 | 0,02 | 4,32E-01 |
| Peg3 | *18616* | -1,77 | 5,07E-11 | -1,52 | 1,20E-08 | 0,48 | 1,00E+00 |
| Suox | *211389* | 1,77 | 1,63E-05 | 1,76 | 8,02E-10 | 0,89 | 1,00E+00 |
| Glipr1 | *73690* | -2,11 | 8,40E-03 | -2,07 | 1,47E-03 | 0,98 | 1,00E+00 |
| Cdh2 | *12558* | -1,55 | 1,64E-08 | -2,05 | 2,28E-13 | 0,11 | 1,00E+00 |
| Osgin1 | *71839* | -1,68 | 3,40E-03 | -1,50 | 6,51E-03 | 0,5 | 1,00E+00 |
| Pcsk9 | *100102* | 1,81 | 2,13E-04 | 3,02 | 8,91E-09 | 0 | 8,94E-02 |
| St3gal6 | *54613* | 2,37 | 6,04E-08 | 2,76 | 4,63E-10 | 0,25 | 1,00E+00 |
| Plekhf1 | *72287* | 1,61 | 2,49E-06 | 1,52 | 8,95E-04 | 0,7 | 1,00E+00 |
| Dusp1 | *19252* | -2,26 | 4,46E-03 | -2,51 | 1,09E-03 | 0,75 | 1,00E+00 |
| Serpinb1a | *66222* | 3,33 | 1,23E-06 | 5,05 | 2,02E-14 | 0 | 1,76E-02 |
| Pacrg | *69310* | 2,88 | 6,31E-05 | 4,36 | 5,38E-36 | 0 | 4,22E-03 |
| Gm36117 | *102639918* | 1,64 | 1,34E-04 | 2,56 | 2,33E-05 | 0,01 | 3,50E-01 |
| Ifi203 | *15950* | 2,74 | 1,31E-03 | 4,51 | 1,18E-07 | 0 | 9,47E-02 |
| Cryab | *12955* | -2,42 | 2,91E-06 | -4,03 | 9,02E-14 | 0,01 | 3,48E-01 |
| Deptor | *97998* | 2,34 | 4,19E-12 | 3,04 | 4,58E-05 | 0,12 | 1,00E+00 |
| Shf | *435684* | 1,51 | 1,48E-03 | 2,72 | 2,05E-03 | 0,03 | 6,15E-01 |
| Nes | *18008* | -2,07 | 2,18E-14 | -2,18 | 7,73E-16 | 0,81 | 1,00E+00 |
| Bicc1 | *83675* | -2,15 | 7,14E-16 | -2,98 | 8,07E-28 | 0,17 | 1,00E+00 |
| Dglucy | *217830* | -1,85 | 2,81E-09 | -2,10 | 1,61E-14 | 0,43 | 1,00E+00 |
| Aif1l | *108897* | -1,51 | 2,86E-05 | -2,38 | 7,55E-11 | 0,08 | 9,46E-01 |
| Aldh1a1 | *11668* | -2,99 | 4,16E-17 | -2,71 | 2,13E-22 | 0,78 | 1,00E+00 |
| Arl14 | *71619* | 2,50 | 5,33E-07 | 3,30 | 4,95E-17 | 0,03 | 6,68E-01 |
| Ifi202b | *26388* | 1,66 | 6,47E-05 | 2,21 | 1,32E-10 | 0,13 | 1,00E+00 |
| Csf1 | *12977* | -1,93 | 4,01E-03 | -2,83 | 1,97E-05 | 0,07 | 9,32E-01 |
| Cdc42bpg | *240505* | -1,79 | 3,01E-06 | -1,73 | 3,29E-07 | 0,9 | 1,00E+00 |
| Egfr | *13649* | -1,99 | 2,71E-13 | -2,86 | 4,62E-26 | 0,02 | 4,71E-01 |
| Bhlhb9 | *70237* | 2,04 | 1,17E-10 | 1,93 | 4,01E-10 | 0,71 | 1,00E+00 |
| Bcas1 | *76960* | 2,80 | 1,60E-10 | 3,01 | 7,32E-06 | 0,54 | 1,00E+00 |
| Dmbt1 | *12945* | 2,95 | 2,65E-08 | 4,43 | 1,59E-14 | 0 | 3,42E-03 |
| Nrg1 | *211323* | -1,71 | 4,33E-09 | -2,09 | 2,68E-14 | 0,36 | 1,00E+00 |
| Dcxr | *67880* | -2,14 | 9,29E-07 | -2,26 | 8,62E-10 | 0,84 | 1,00E+00 |
| Ak1 | *11636* | -2,20 | 1,89E-14 | -3,13 | 1,38E-28 | 0,19 | 1,00E+00 |
| Cdhr5 | *72040* | 2,20 | 7,48E-05 | 3,06 | 1,34E-16 | 0,02 | 4,98E-01 |
| Pdk1 | *228026* | 1,87 | 2,73E-03 | 2,78 | 3,43E-03 | 0,11 | 1,00E+00 |
| Slc4a11 | *269356* | -1,80 | 9,82E-04 | -2,16 | 2,62E-09 | 0,53 | 1,00E+00 |
| Fcho1 | *74015* | 2,31 | 2,07E-05 | 2,54 | 2,69E-08 | 0,53 | 1,00E+00 |
| Akap5 | *238276* | -2,90 | 4,38E-07 | -4,86 | 1,66E-15 | 0,03 | 5,65E-01 |
| Kcnh2 | *16511* | 3,17 | 6,31E-04 | 3,62 | 8,01E-03 | 0,49 | 1,00E+00 |
| Trim7 | *94089* | -1,55 | 2,82E-03 | -1,94 | 1,30E-08 | 0,58 | 1,00E+00 |
| Serpinb6b | *20708* | -1,72 | 1,91E-03 | -3,38 | 1,57E-14 | 0,08 | 9,57E-01 |
| Adgrg6 | *215798* | -1,83 | 1,26E-10 | -2,22 | 8,59E-16 | 0,4 | 1,00E+00 |
| Gsn | *227753* | -2,14 | 4,10E-11 | -3,11 | 2,73E-22 | 0,01 | 2,95E-01 |
| Gstm1 | *14862* | -1,80 | 7,27E-02 | -1,91 | 1,04E-02 | 0,89 | 1,00E+00 |
| Arid5a | *214855* | 1,94 | 4,53E-06 | 2,81 | 7,42E-15 | 0,01 | 3,38E-01 |
| Mst1r | *19882* | 2,73 | 6,72E-13 | 2,55 | 1,61E-06 | 0,63 | 1,00E+00 |
| Slc16a6 | *104681* | 1,88 | 3,02E-05 | 2,15 | 8,10E-08 | 0,4 | 1,00E+00 |
| Bscl2 | *14705* | -1,51 | 7,47E-06 | -2,35 | 8,87E-14 | 0,03 | 5,65E-01 |
| Fgd3 | *30938* | -2,53 | 2,38E-03 | -3,41 | 2,28E-05 | 0,07 | 9,23E-01 |
| Gadd45b | *17873* | -1,74 | 2,99E-07 | -3,15 | 1,47E-20 | 0 | 1,74E-01 |
| Hoxa3 | *15400* | 1,50 | 3,10E-03 | 1,83 | 5,16E-06 | 0,38 | 1,00E+00 |
| Pfkp | *56421* | 1,58 | 1,79E-04 | 2,37 | 9,77E-04 | 0,05 | 8,03E-01 |
| Tnfrsf1b | *21938* | 1,73 | 4,79E-04 | 3,53 | 1,24E-06 | 0 | 3,66E-02 |
| Foxq1 | *15220* | -1,79 | 2,82E-07 | -2,15 | 7,55E-11 | 0,42 | 1,00E+00 |
| Slco3a1 | *108116* | -1,72 | 3,89E-09 | -2,42 | 9,27E-17 | 0,04 | 7,36E-01 |
| Tns1 | *21961* | -2,55 | 3,88E-08 | -2,83 | 3,00E-10 | 0,67 | 1,00E+00 |
| Psca | *72373* | -2,53 | 1,18E-02 | -3,67 | 8,04E-05 | 0,11 | 1,00E+00 |
| Fn1 | *14268* | -3,57 | 1,19E-05 | -3,87 | 2,12E-06 | 0,77 | 1,00E+00 |
| Plxna4 | *243743* | 3,05 | 1,75E-04 | 4,22 | 3,28E-09 | 0,01 | 2,63E-01 |
| Rcor2 | *104383* | 1,56 | 6,11E-03 | 1,69 | 2,68E-04 | 0,77 | 1,00E+00 |
| Rassf9 | *237504* | -1,63 | 2,60E-03 | -1,56 | 2,73E-03 | 0,88 | 1,00E+00 |
| Glipr2 | *384009* | -2,71 | 2,59E-12 | -3,80 | 4,94E-21 | 0,03 | 6,54E-01 |
| Serpinb9b | *20706* | -2,75 | 2,20E-02 | -5,02 | 2,88E-06 | 0,16 | 1,00E+00 |
| Kank1 | *107351* | -2,96 | 1,09E-13 | -5,26 | 2,76E-32 | 0,01 | 4,08E-01 |
| Hoxb3 | *15410* | -3,35 | 2,97E-11 | -2,93 | 6,14E-11 | 0,59 | 1,00E+00 |

| | | | | | | | |
|---|---|---|---|---|---|---|---|
| Hoxb5 | 15413 | -4,81 | 4,62E-11 | -3,89 | 3,10E-10 | 0,54 | 1,00E+00 |
| Hoxb8 | 15416 | -3,43 | 2,34E-10 | -1,89 | 5,10E-05 | 0,02 | 4,51E-01 |
| Mtus1 | 102103 | -1,99 | 2,16E-08 | -3,44 | 1,11E-22 | 0,05 | 7,65E-01 |
| Ces2g | 72361 | -4,44 | 1,55E-07 | -5,99 | 1,94E-11 | 0,17 | 1,00E+00 |
| Serpinb8 | 20725 | -2,82 | 2,68E-07 | -5,47 | 3,28E-16 | 0,01 | 3,38E-01 |
| Fam149a | 212326 | -3,05 | 2,76E-05 | -5,02 | 1,49E-11 | 0,11 | 1,00E+00 |
| P3h2 | 210530 | -1,63 | 7,72E-05 | -3,20 | 2,29E-11 | 0 | 5,49E-02 |
| Zfp764 | 233893 | -2,77 | 1,59E-04 | -3,77 | 1,19E-08 | 0,32 | 1,00E+00 |
| H60b | 667281 | -3,28 | 1,76E-04 | -4,68 | 1,27E-07 | 0,43 | 1,00E+00 |
| Sorl1 | 20660 | -1,73 | 1,84E-03 | -3,07 | 2,73E-08 | 0 | 4,26E-02 |
| Ccbe1 | 320924 | -1,83 | 5,45E-02 | -3,41 | 1,51E-09 | 0,34 | 1,00E+00 |
| Htatip2 | 53415 | -4,78 | 7,44E-11 | -5,95 | 1,66E-15 | 0,62 | 1,00E+00 |
| Ces2e | 234673 | -5,68 | 1,02E-06 | -6,98 | 6,90E-09 | 0,51 | 1,00E+00 |
| Cutal | 77996 | -2,32 | 7,89E-06 | -1,85 | 5,15E-05 | 0,6 | 1,00E+00 |
| Gm9949 | 225609 | -3,19 | 1,87E-05 | -5,20 | 7,90E-12 | 0,16 | 1,00E+00 |
| Afap1l2 | 226250 | -3,98 | 3,61E-05 | -4,89 | 2,45E-07 | 0,14 | 1,00E+00 |
| Gsta3 | 14859 | -7,08 | 8,13E-05 | -8,26 | 7,37E-06 | 0 | 1,00E+00 |
| Tgfb2 | 21808 | -1,71 | 7,88E-05 | -3,69 | 6,54E-23 | 0,07 | 9,16E-01 |
| Notch3 | 18131 | -3,37 | 1,88E-04 | -4,14 | 9,88E-07 | 0,5 | 1,00E+00 |
| Samd9l | 209086 | -2,67 | 3,56E-03 | -3,57 | 2,09E-05 | 0,26 | 1,00E+00 |
| Dusp8 | 18218 | -1,52 | 3,11E-02 | -2,02 | 1,30E-03 | 0,27 | 1,00E+00 |
| Dkk2 | 56811 | -1,95 | 5,67E-02 | -3,54 | 6,67E-05 | 0,06 | 8,83E-01 |
| Wt1 | 22431 | -3,39 | 5,56E-15 | -6,26 | 1,54E-35 | 0,05 | 8,01E-01 |
| Cd59a | 12509 | -3,40 | 1,17E-10 | -5,92 | 5,37E-24 | 0,16 | 1,00E+00 |
| Mmp2 | 17390 | -2,43 | 3,46E-09 | -3,85 | 1,50E-21 | 0,06 | 8,53E-01 |
| Smarca2 | 67155 | -2,56 | 3,46E-08 | -3,47 | 5,08E-15 | 0,28 | 1,00E+00 |
| Serpina1a | 20700 | -4,41 | 1,61E-05 | -5,35 | 1,17E-07 | 0 | 1,00E+00 |
| Ctla2b | 13025 | -2,63 | 7,75E-05 | -7,35 | 5,86E-17 | 0 | 1,57E-01 |
| Pla1a | 85031 | -2,04 | 2,36E-03 | -2,23 | 1,45E-07 | 0,85 | 1,00E+00 |
| Nid2 | 18074 | -1,90 | 1,19E-02 | -4,25 | 3,86E-08 | 0 | 1,26E-01 |
| Nnat | 18111 | -3,31 | 1,85E-02 | -4,91 | 3,15E-04 | 0,44 | 1,00E+00 |
| Shank2 | 210274 | -2,67 | 4,92E-17 | -3,33 | 7,85E-27 | 0,27 | 1,00E+00 |
| Gria3 | 53623 | -5,65 | 7,64E-47 | -9,67 | 1,20E-71 | 0 | 2,12E-01 |
| Akr1c14 | 105387 | -4,51 | 1,37E-23 | -8,75 | 2,06E-43 | 0,07 | 9,12E-01 |
| Mapkapk3 | 102626 | -1,85 | 1,27E-05 | -1,74 | 9,88E-07 | 0,91 | 1,00E+00 |
| Gdpd5 | 233552 | -1,54 | 1,71E-05 | -1,92 | 4,37E-09 | 0,41 | 1,00E+00 |
| Heg1 | 77446 | -2,04 | 1,94E-04 | -1,89 | 5,97E-04 | 0,76 | 1,00E+00 |
| Spns2 | 216892 | -1,65 | 2,57E-04 | -2,59 | 6,48E-08 | 0,03 | 6,01E-01 |
| Serpinb9 | 20723 | -1,71 | 5,69E-04 | -3,27 | 1,62E-21 | 0,16 | 1,00E+00 |
| Rnf130 | 59044 | -3,83 | 1,07E-03 | -6,65 | 3,31E-07 | 0,07 | 9,09E-01 |
| Rab38 | 72433 | -4,45 | 3,76E-02 | -5,48 | 4,75E-03 | 0,59 | 1,00E+00 |
| Ccn4 | 22402 | -1,90 | 3,56E-02 | -5,04 | 3,74E-11 | 0,04 | 7,37E-01 |
| Nectin1 | 58235 | -2,14 | 1,45E-11 | -2,95 | 4,20E-21 | 0,04 | 7,16E-01 |
| Mab21l4 | 71874 | -2,49 | 3,09E-11 | -4,16 | 3,68E-28 | 0,1 | 1,00E+00 |
| Pik3ap1 | 83490 | -2,12 | 1,45E-09 | -3,31 | 8,85E-24 | 0,05 | 7,65E-01 |
| Ckmt1 | 12716 | -3,03 | 2,22E-05 | -4,51 | 6,43E-10 | 0,15 | 1,00E+00 |
| Cox6b2 | 333182 | -3,34 | 3,03E-03 | -5,08 | 6,53E-06 | 0,42 | 1,00E+00 |
| Tmem14a | 75712 | -1,70 | 2,89E-03 | -2,98 | 1,28E-09 | 0,1 | 1,00E+00 |
| Def6 | 23853 | -2,41 | 3,77E-03 | -2,95 | 2,29E-04 | 0,34 | 1,00E+00 |
| Sytl1 | 269589 | -3,71 | 5,84E-03 | -4,71 | 3,02E-04 | 0,31 | 1,00E+00 |
| Siglecg | 243958 | -3,91 | 5,77E-03 | -6,30 | 9,92E-06 | 0,01 | 3,75E-01 |
| Tap1 | 21354 | -3,06 | 3,82E-16 | -3,96 | 2,05E-26 | 0,23 | 1,00E+00 |
| Tbc1d8 | 54610 | -2,32 | 6,68E-12 | -3,35 | 4,28E-24 | 0,05 | 7,66E-01 |
| Ctla2a | 13024 | -2,82 | 2,53E-08 | -5,17 | 1,37E-21 | 0,01 | 2,62E-01 |
| Ly6e | 17069 | -3,96 | 9,60E-05 | -5,84 | 1,38E-08 | 0,2 | 1,00E+00 |
| 9230114K14Rik | 414108 | -1,63 | 9,00E-05 | -2,48 | 2,53E-11 | 0,06 | 8,97E-01 |
| Hoxb7 | 15415 | -4,78 | 2,42E-20 | -5,08 | 1,16E-24 | 0,79 | 1,00E+00 |
| Hoxb6 | 15414 | -3,91 | 2,62E-16 | -3,53 | 3,96E-16 | 0,64 | 1,00E+00 |
| Serpina1b | 20701 | -5,09 | 7,17E-13 | -5,18 | 1,05E-13 | 0,97 | 1,00E+00 |
| Wnt9a | 216795 | -3,56 | 7,22E-10 | -6,78 | 4,63E-23 | 0 | 1,53E-01 |
| Mcam | 84004 | -4,11 | 4,04E-07 | -5,67 | 3,63E-11 | 0,05 | 8,01E-01 |
| Cd109 | 235505 | -1,68 | 3,59E-04 | -3,17 | 1,63E-10 | 0,01 | 2,28E-01 |
| Pros1 | 19128 | -1,96 | 9,33E-03 | -2,45 | 4,32E-04 | 0,23 | 1,00E+00 |
| Inka2 | 109050 | -1,68 | 1,99E-07 | -2,77 | 5,43E-20 | 0,02 | 4,52E-01 |
| Dlc1 | 50768 | -4,75 | 7,63E-05 | -8,52 | 3,05E-10 | 0,09 | 1,00E+00 |
| Gstm2 | 14863 | -1,96 | 6,77E-03 | -2,91 | 3,93E-06 | 0,24 | 1,00E+00 |
| Fas | 14102 | -3,94 | 2,55E-17 | -5,91 | 5,90E-32 | 0,26 | 1,00E+00 |
| Gm38426 | 100503676 | -2,14 | 6,67E-05 | -2,35 | 1,96E-06 | 0,7 | 1,00E+00 |
| Celf5 | 319586 | -2,96 | 2,45E-03 | -3,71 | 2,89E-05 | 0,41 | 1,00E+00 |
| Bmyc | 107771 | -2,59 | 3,77E-03 | -5,34 | 2,28E-07 | 0,01 | 2,74E-01 |
| Tubb4a | 22153 | -4,07 | 1,30E-02 | -3,97 | 7,68E-03 | 0,85 | 1,00E+00 |
| Pclaf | 68026 | -1,73 | 1,86E-02 | -3,90 | 1,69E-09 | 0,04 | 7,40E-01 |
| Scx | 20289 | -2,28 | 4,96E-08 | -4,29 | 1,67E-24 | 0,15 | 1,00E+00 |
| Creb3l1 | 26427 | -1,90 | 5,46E-02 | -2,86 | 1,50E-05 | 0,44 | 1,00E+00 |
| Ano3 | 228432 | -2,86 | 3,40E-16 | -4,72 | 8,69E-48 | 0,09 | 1,00E+00 |
| Ddit4l | 73284 | -3,35 | 5,16E-24 | -6,24 | 1,98E-60 | 0 | 1,99E-01 |
| Arhgef26 | 622434 | -1,53 | 2,97E-03 | -5,32 | 3,60E-38 | 0 | 1,04E-01 |
| Foxj1 | 15223 | -2,87 | 6,40E-21 | -4,84 | 2,96E-50 | 0,09 | 1,00E+00 |
| 1700007K13Rik | 69327 | -4,53 | 4,18E-08 | -6,63 | 3,42E-13 | 0,08 | 9,47E-01 |
| Cdh6 | 12563 | -2,48 | 4,04E-03 | -3,71 | 1,03E-05 | 0,1 | 1,00E+00 |
| Ndn | 17984 | -6,93 | 2,59E-06 | -9,82 | 2,56E-09 | 0,19 | 1,00E+00 |
| Igfbp7 | 29817 | -4,02 | 7,18E-44 | -8,11 | 2,39E-111 | 0 | 9,92E-02 |

**Supplementary Table 2: gene expression modified only in Ras$^{HIGH}$ cells (Figure 1E middle panel)**

| n=65 | | Ras Low vs WT | | Ras High vs WT | | Ras High vs Ras Low | |
|---|---|---|---|---|---|---|---|
| Gene symbol | Gene id | log2FoldChange | padjExactTest | log2FoldChange | padjExactTest | log2FoldChange | padjExactTest |
| **Prl2c2** | *18811* | 2,68 | 7,54E-01 | 6,57 | 1,80E-19 | 3,88 | 1,88E-08 |
| **Anxa13** | *69787* | 0,38 | 9,24E-01 | 2,52 | 4,21E-21 | 2,14 | 1,40E-07 |
| **Apoc2** | *11813* | 1,40 | 9,63E-01 | 5,46 | 5,51E-12 | 4,05 | 5,14E-07 |
| **Prl2c5** | *107849* | 2,80 | 7,34E-01 | 6,00 | 5,40E-43 | 3,19 | 8,57E-12 |
| **Il1rn** | *16181* | 1,30 | 9,13E-01 | 5,57 | 5,12E-47 | 4,25 | 6,47E-28 |
| **Sord** | *20322* | 1,42 | 1,14E-01 | 3,32 | 4,39E-09 | 1,89 | 2,26E-03 |
| **Arap3** | *106952* | 2,42 | 1,21E-01 | 4,46 | 1,86E-54 | 2,03 | 3,47E-07 |
| **Mrc1** | *17533* | 1,69 | 2,03E-01 | 3,33 | 2,18E-25 | 1,63 | 4,38E-04 |
| **Glrp1** | *14659* | 1,62 | 1,79E-01 | 3,31 | 4,21E-13 | 1,69 | 3,70E-03 |
| **Pla2g7** | *27226* | 1,53 | 1,31E-01 | 3,63 | 1,59E-16 | 2,10 | 6,71E-06 |
| **Kcnn4** | *16534* | 2,57 | 2,70E-01 | 5,27 | 8,53E-19 | 2,69 | 3,61E-06 |
| **Mndal** | *100040462* | 2,28 | 2,74E-02 | 4,35 | 4,34E-10 | 2,06 | 5,53E-03 |
| **Tm4sf1** | *17112* | 2,15 | 3,35E-02 | 6,32 | 1,66E-15 | 4,17 | 1,79E-07 |
| **Rhox5** | *18617* | 2,24 | 2,04E-02 | 3,96 | 2,16E-15 | 1,72 | 5,33E-03 |
| **Sigirr** | *24058* | 0,90 | 9,53E-01 | 3,30 | 7,25E-13 | 2,39 | 9,59E-04 |
| **Ugt1a7c** | *394432* | -1,56 | 7,69E-01 | 1,90 | 6,56E-03 | 3,45 | 7,18E-08 |
| **Syt7** | *54525* | 2,20 | 1,96E-01 | 4,51 | 1,92E-48 | 2,30 | 1,88E-08 |
| **Dab2** | *13132* | -0,48 | 3,33E-01 | -2,23 | 3,80E-14 | -1,75 | 1,80E-05 |
| **Selp** | *20344* | 3,30 | 3,84E-02 | 5,14 | 2,30E-43 | 1,84 | 1,07E-05 |
| **Usp43** | *216835* | 1,26 | 2,57E-02 | 3,02 | 8,41E-21 | 1,76 | 1,17E-04 |
| **Inpp5j** | *170835* | 0,75 | 6,85E-01 | 2,93 | 1,35E-13 | 2,17 | 4,60E-07 |
| **Tmem173** | *72512* | -0,40 | 8,50E-01 | 1,51 | 2,20E-03 | 1,90 | 1,75E-03 |
| **Havcr2** | *171285* | 1,55 | 5,38E-01 | 5,51 | 1,38E-16 | 3,95 | 2,79E-10 |
| **Steap3** | *68428* | 1,69 | 9,16E-02 | 4,51 | 3,25E-07 | 2,81 | 3,08E-03 |
| **Csn3** | *12994* | 0,03 | 1,00E+00 | 3,57 | 1,10E-03 | 3,53 | 9,32E-03 |
| **Kif5c** | *16574* | -0,40 | 8,36E-01 | 2,75 | 8,93E-12 | 3,14 | 7,52E-14 |
| **Map3k6** | *53608* | 0,21 | 1,00E+00 | 1,98 | 8,61E-09 | 1,76 | 3,25E-04 |
| **Rgl1** | *19731* | -0,40 | 8,25E-01 | 1,94 | 1,70E-06 | 2,34 | 2,29E-07 |
| **Slco4a1** | *108115* | -2,68 | 2,06E-02 | 1,52 | 7,33E-03 | 4,19 | 6,41E-10 |
| **Spaca9** | *69987* | 0,11 | 1,00E+00 | 2,80 | 1,73E-04 | 2,68 | 5,53E-03 |
| **Tmem252** | *226040* | -0,62 | 9,74E-01 | 3,94 | 2,59E-08 | 4,56 | 3,67E-08 |
| **Trim14** | *74735* | 0,46 | 7,36E-01 | 2,30 | 4,87E-07 | 1,84 | 3,71E-04 |
| **Epdr1** | *105298* | -0,84 | 2,24E-02 | -2,46 | 6,01E-17 | -1,62 | 6,91E-04 |
| **Clrn3** | *212070* | 0,64 | 8,81E-01 | 2,43 | 5,45E-09 | 1,79 | 1,72E-03 |
| **Cyth4** | *72318* | -0,23 | 1,00E+00 | 1,61 | 2,62E-05 | 1,83 | 1,01E-03 |
| **Naip1** | *17940* | 0,56 | 8,48E-01 | 3,27 | 2,33E-08 | 2,71 | 1,72E-05 |
| **Qrfp** | *227717* | -0,23 | 9,93E-01 | 2,10 | 2,81E-03 | 2,33 | 7,75E-03 |
| **Sh3bp2** | *24055* | -0,09 | 1,00E+00 | 1,59 | 1,73E-03 | 1,68 | 4,02E-03 |
| **Hs6st2** | *50786* | -0,94 | 3,58E-03 | -3,79 | 1,10E-39 | -2,84 | 1,13E-12 |
| **Cd53** | *12508* | 2,52 | 7,52E-02 | 5,45 | 3,21E-09 | 2,92 | 1,32E-03 |
| **Ceacam1** | *26365* | 1,98 | 7,19E-01 | 5,52 | 2,13E-07 | 3,53 | 2,25E-03 |
| **Sec16b** | *89867* | 0,10 | 9,99E-01 | 2,46 | 5,00E-09 | 2,36 | 9,91E-07 |
| **Slc14a1** | *108052* | 2,47 | 8,55E-01 | 5,27 | 1,25E-21 | 2,79 | 1,34E-06 |
| **Tnxb** | *81877* | 1,97 | 1,86E-01 | 3,62 | 2,49E-26 | 1,65 | 8,74E-04 |
| **Map2** | *17756* | -0,07 | 1,00E+00 | -2,60 | 5,94E-04 | -2,53 | 4,07E-07 |
| **Shroom3** | *27428* | 0,22 | 9,56E-01 | -1,72 | 4,47E-08 | -1,94 | 6,60E-03 |
| **Them6** | *223626* | -1,31 | 1,08E-03 | -2,99 | 2,01E-16 | -1,68 | 1,19E-02 |
| **9330188P03Rik** | *380930* | 3,10 | 8,04E-01 | 6,74 | 5,79E-11 | 3,64 | 8,60E-05 |
| **Cdh5** | *12562* | 4,04 | 5,34E-01 | 6,87 | 1,02E-20 | 2,82 | 2,95E-05 |
| **Tlr4** | *21898* | 0,97 | 2,55E-01 | 2,96 | 2,21E-09 | 1,98 | 4,42E-05 |
| **Pla2g16** | *225845* | -1,35 | 4,76E-02 | -3,44 | 4,05E-07 | -2,09 | 4,09E-03 |
| **Lrp2** | *14725* | -1,30 | 3,90E-01 | -4,23 | 3,25E-04 | -2,92 | 5,92E-02 |
| **Pcsk5** | *18552* | -2,53 | 5,80E-02 | -5,90 | 1,35E-05 | -3,37 | 3,54E-02 |
| **Cep126** | *234915* | -0,75 | 2,81E-01 | -2,41 | 4,91E-09 | -1,66 | 1,82E-02 |
| **Zfp185** | *22673* | -0,85 | 1,43E-01 | -2,45 | 2,21E-09 | -1,60 | 3,62E-02 |
| **Fgfr3** | *14184* | -1,37 | 9,86E-04 | -3,89 | 1,65E-18 | -2,51 | 3,32E-03 |
| **Upk3b** | *100647* | -2,20 | 7,06E-02 | -5,67 | 2,55E-06 | -3,46 | 5,88E-02 |
| **Ssbp2** | *66970* | -0,84 | 1,33E-01 | -3,37 | 4,40E-17 | -2,53 | 5,50E-02 |
| **Pkia** | *18767* | -1,25 | 4,79E-04 | -3,46 | 1,26E-24 | -2,21 | 1,72E-05 |
| **Epb41l3** | *13823* | -1,82 | 3,52E-01 | -5,45 | 1,13E-03 | -3,63 | 4,40E-09 |
| **Mpzl2** | *14012* | -0,39 | 7,42E-01 | -2,62 | 6,99E-06 | -2,23 | 1,11E-06 |
| **Npr2** | *230103* | -0,82 | 1,66E-01 | -3,21 | 5,52E-18 | -2,40 | 2,85E-03 |
| **Fgf18** | *14172* | -0,81 | 3,04E-01 | -3,87 | 4,64E-17 | -3,05 | 5,68E-02 |
| **Uchl1** | *22223* | -0,95 | 5,12E-01 | -3,57 | 8,49E-05 | -2,61 | 9,20E-03 |
| **Nuak1** | *77976* | -2,44 | 1,08E-01 | -3,95 | 3,42E-03 | -1,52 | 1,44E-02 |

**Supplementary Table 3: genes for which expression correlated with intensity of Ras/MAPK signalling (Figure 1E right pan**

genes n = 15

| | | Ras Low vs WT | | Ras High vs WT | | Ras High vs Ras Low | |
|---|---|---|---|---|---|---|---|
| Gene symbol | Gene id | log2FoldChange | padjExactTest | log2FoldChange | padjExactTest | log2FoldChange | padjExactTest |
| **AI467606** | *101602* | 2,43 | 1,95E-05 | 4,24 | 1,57E-25 | 1,80 | 6,28E-05 |
| **Aim2** | *383619* | 1,79 | 5,59E-03 | 3,97 | 2,60E-36 | 2,18 | 3,47E-07 |
| **Dynap** | *75577* | 5,84 | 4,86E-03 | 7,92 | 2,35E-97 | 2,08 | 3,43E-07 |
| **Htra3** | *78558* | 1,86 | 1,49E-03 | 3,74 | 8,65E-09 | 1,87 | 5,89E-03 |
| **Itgb7** | *16421* | 5,26 | 3,29E-04 | 6,92 | 1,15E-47 | 1,66 | 3,16E-03 |
| **Tspan13** | *66109* | 1,93 | 8,38E-04 | 3,88 | 7,00E-10 | 1,95 | 2,28E-03 |
| **Ppp2r2b** | *72930* | -2,89 | 2,31E-20 | -8,11 | 8,07E-79 | -5,23 | 8,11E-03 |
| **Cbr1** | *12408* | -2,44 | 9,54E-03 | -5,23 | 2,47E-07 | -2,80 | 4,13E-03 |
| **Pmp22** | *18858* | -2,35 | 3,06E-03 | -4,56 | 5,31E-08 | -2,21 | 7,67E-03 |
| **Ptp4a3** | *19245* | -4,01 | 1,85E-41 | -8,04 | 4,59E-103 | -4,03 | 4,29E-02 |
| **Pmaip1** | *58801* | -2,16 | 1,09E-13 | -4,45 | 1,01E-47 | -2,29 | 4,12E-07 |
| **Thbs1** | *21825* | -3,56 | 5,11E-05 | -5,42 | 3,05E-09 | -1,86 | 4,97E-05 |
| **Akap12** | *83397* | -5,27 | 3,74E-11 | -8,60 | 7,60E-22 | -3,33 | 5,98E-02 |
| **Sulf2** | *72043* | -2,18 | 1,65E-15 | -4,61 | 1,04E-56 | -2,43 | 1,45E-09 |
| **Crip2** | *68337* | -2,45 | 3,10E-05 | -4,07 | 8,05E-11 | -1,62 | 1,63E-04 |

**Supplementary Table 4: genes differentialy expressed in peritoneal tumor cells vs liver tumor cells**

log2 fold change >1; p<0.05

| Gene symbol | Gene ID | log2FoldChange | p-value |
|---|---|---|---|
| Fgg | 99571 | Inf | 1,05E-06 |
| Fga | 14161 | Inf | 3,81E-06 |
| Clec4f | 51811 | Inf | 2,61E-07 |
| Mug1 | 17836 | Inf | 1,04E-05 |
| Serpina3k | 20714 | Inf | 3,42E-04 |
| Clec4g | 75863 | Inf | 3,74E-05 |
| Saa1 | 20208 | Inf | 4,97E-03 |
| C8a | 230558 | Inf | 5,12E-03 |
| Hrg | 94175 | Inf | 5,94E-03 |
| Mat1a | 11720 | Inf | 2,90E-02 |
| Fgl1 | 234199 | 6,51 | 1,39E-03 |
| Glp1r | 19142 | 4,07 | 2,25E-12 |
| Prss12 | 11625 | 3,13 | 6,29E-03 |
| Ahsg | 12705 | 6,18 | 2,10E-06 |
| Cited1 | 12994 | 2,97 | 3,58E-04 |
| Csn3 | 11287 | 2,92 | 2,65E-19 |
| Pzp | 11657 | 2,83 | 4,27E-03 |
| Alb | 233107 | 6,09 | 1,39E-04 |
| Kctd15 | 110135 | 2,66 | 1,02E-02 |
| Fgb | 108832 | 5,65 | 1,90E-05 |
| Tmem74b | 18162 | 2,54 | 4,54E-05 |
| Npr3 | 22295 | 2,45 | 1,40E-04 |
| Cdh23 | 16427 | 2,40 | 4,58E-03 |
| Itih4 | 11803 | 5,54 | 3,57E-05 |
| Aplp1 | 381853 | 2,25 | 1,54E-10 |
| Gipr | 18599 | 2,12 | 3,32E-24 |
| Padi1 | 26365 | 2,04 | 8,66E-11 |
| Ceacam1 | 319875 | 1,91 | 1,54E-06 |
| Tmprss11b | 20408 | 1,91 | 2,08E-07 |
| Sh3gl3 | 338372 | 1,79 | 3,40E-05 |
| Map3k9 | 100503085 | 1,76 | 2,52E-03 |
| Klhl3 | 21916 | 1,71 | 7,81E-09 |
| Tmod1 | 320311 | 1,69 | 1,21E-05 |
| Rnf152 | 14745 | 1,69 | 2,43E-02 |
| Lpar1 | 67749 | 1,66 | 3,89E-17 |
| Mgarp | 20148 | 1,64 | 5,12E-09 |
| Dhrs3 | 15483 | 1,61 | 5,15E-07 |
| Hsd11b1 | 243084 | 1,60 | 3,05E-05 |
| Tmprss11e | 319924 | 1,56 | 2,22E-17 |
| Apba1 | 20667 | 1,55 | 9,39E-09 |
| Sox12 | 70839 | 1,47 | 6,86E-03 |
| P2ry12 | 78560 | 1,37 | 2,31E-06 |
| Adgra2 | 14373 | 1,36 | 5,36E-05 |
| G0s2 | 14360 | 1,34 | 1,28E-06 |
| Fyn | 108078 | 1,33 | 6,46E-14 |
| Olr1 | 226518 | 1,31 | 3,74E-07 |

| | | | |
|---|---|---|---|
| Nmnat2 | 224796 | 1,30 | 6,71E-04 |
| Clic5 | 100039691 | 1,29 | 5,93E-04 |
| Lncenc1 | 16565 | 1,27 | 6,87E-05 |
| Kif21b | 94227 | 1,27 | 9,95E-10 |
| Pi15 | 226040 | 1,26 | 4,90E-04 |
| Tmem252 | 15118 | 1,25 | 7,88E-08 |
| Has3 | 20344 | 1,15 | 5,31E-06 |
| Selp | 16372 | 1,14 | 1,42E-05 |
| Irx2 | 14064 | 1,12 | 1,11E-04 |
| F2rl2 | 245884 | 1,11 | 9,74E-04 |
| Fam71f2 | 67216 | 1,09 | 9,05E-04 |
| Mboat2 | 242819 | 1,08 | 3,84E-07 |
| Rundc3b | 67547 | 1,05 | 2,63E-03 |
| Slc39a8 | 16780 | 1,04 | 5,82E-03 |
| Lamb3 | 14652 | 1,02 | 7,02E-09 |
| Pcolce | 18542 | -1,01 | 7,15E-06 |
| Clec2d | 93694 | -1,02 | 1,01E-04 |
| Slc22a18 | 18400 | -1,03 | 3,14E-06 |
| Daglb | 231871 | -1,03 | 9,31E-04 |
| Cercam | 99151 | -1,04 | 1,09E-03 |
| Cgn | 70737 | -1,04 | 1,14E-03 |
| Cfb | 14962 | -1,05 | 9,75E-05 |
| Cyp27a1 | 104086 | -1,08 | 2,15E-05 |
| Foxj1 | 15223 | -1,08 | 2,04E-03 |
| Apoc2 | 11813 | -1,09 | 3,92E-03 |
| Gstm1 | 14862 | -1,09 | 1,43E-03 |
| Adamtsl5 | 66548 | -1,10 | 2,98E-06 |
| Sparc | 20692 | -1,10 | 8,87E-05 |
| Ildr2 | 100039795 | -1,11 | 5,45E-03 |
| AW112010 | 107350 | -1,12 | 4,92E-03 |
| Gcnt3 | 72077 | -1,12 | 5,05E-08 |
| Als2cl | 235633 | -1,13 | 5,30E-12 |
| Camsap3 | 69697 | -1,14 | 1,98E-05 |
| Ltbp2 | 16997 | -1,17 | 7,64E-04 |
| Psrc1 | 56742 | -1,19 | 2,19E-08 |
| C2 | 12263 | -1,21 | 8,61E-04 |
| Cbs | 12411 | -1,24 | 9,02E-03 |
| Uba7 | 74153 | -1,25 | 7,85E-04 |
| Cidec | 14311 | -1,39 | 7,22E-03 |
| Lgals4 | 16855 | -1,40 | 7,44E-03 |
| Apob | 238055 | -1,45 | 6,72E-06 |
| Sema3f | 20350 | -1,49 | 5,34E-08 |
| Cideb | 12684 | -1,52 | 1,98E-05 |
| Plcxd3 | 239318 | -1,66 | 7,08E-04 |
| Bcas1 | 76960 | -1,76 | 1,27E-03 |
| Syt8 | 55925 | -1,77 | 1,73E-16 |
| Gjb1 | 14618 | -1,78 | 1,02E-03 |
| Trim71 | 636931 | -1,80 | 7,74E-03 |
| Mamld1 | 333639 | -1,80 | 3,61E-03 |
| Col5a1 | 12831 | -1,83 | 1,77E-02 |
| Igfbp6 | 16012 | -1,85 | 2,97E-02 |
| Tnc | 21923 | -1,94 | 8,74E-03 |

| | | | |
|---|---|---|---|
| Cdhr5 | 72040 | -1,96 | 7,91E-08 |
| Tnni2 | 21953 | -1,99 | 7,60E-04 |
| Pla1a | 85031 | -2,01 | 1,82E-03 |
| Col15a1 | 12819 | -2,10 | 2,17E-02 |
| Clrn3 | 212070 | -2,15 | 4,13E-06 |
| Col5a3 | 53867 | -2,16 | 1,32E-02 |
| Exoc3l4 | 74190 | -2,17 | 2,70E-04 |
| Tnfaip6 | 21930 | -2,22 | 3,00E-02 |
| Misp | 78906 | -2,28 | 9,05E-41 |
| Cspg4 | 121021 | -2,41 | 4,11E-03 |
| Mrc2 | 17534 | -2,41 | 2,52E-02 |
| Spon1 | 233744 | -2,60 | 1,78E-02 |
| Pdgfrb | 18596 | -2,67 | 2,56E-02 |
| Col16a1 | 107581 | -2,69 | 2,63E-02 |
| Acta2 | 11475 | -2,75 | 1,80E-03 |
| Serpine2 | 20720 | -2,83 | 4,63E-02 |
| Gas1 | 14451 | -2,83 | 1,22E-02 |
| Has1 | 15116 | -3,05 | 1,17E-02 |
| Col1a1 | 12842 | -3,07 | 1,47E-02 |
| Mmp2 | 17390 | -3,08 | 2,79E-02 |
| Serpinf1 | 20317 | -3,09 | 4,06E-02 |
| Olfml3 | 99543 | -3,15 | 1,23E-02 |
| Col1a2 | 12843 | -3,24 | 2,24E-02 |
| Col5a2 | 12832 | -3,31 | 3,24E-02 |
| Srpx2 | 68792 | -3,47 | 3,63E-02 |
| Smoc2 | 64074 | -3,65 | 9,59E-03 |
| Col12a1 | 12816 | -3,76 | 9,76E-08 |
| Ppef1 | 237178 | -3,83 | 4,72E-02 |
| Sod3 | 20657 | -3,89 | 1,66E-02 |
| Loxl1 | 16949 | -3,97 | 2,09E-02 |
| Thy1 | 21838 | -4,10 | 2,29E-03 |
| Cthrc1 | 68588 | -4,12 | 1,40E-03 |
| Fbn1 | 14118 | -4,21 | 4,00E-02 |
| Cxcl12 | 20315 | -4,23 | 4,13E-02 |
| Col3a1 | 12825 | -4,24 | 4,22E-02 |
| Col8a1 | 12837 | -4,41 | 2,48E-03 |
| Fndc1 | 68655 | -4,75 | 3,01E-02 |
| Sfrp1 | 20377 | -4,77 | 2,62E-02 |
| Thbs2 | 21826 | -4,87 | 4,67E-04 |
| Mfap2 | 17150 | -4,96 | 5,56E-04 |
| Eln | 13717 | -4,98 | 4,59E-02 |
| Postn | 50706 | -5,32 | 9,48E-03 |
| Cilp | 214425 | -5,49 | 1,72E-02 |
| Mfap5 | 50530 | -5,60 | 1,45E-02 |
| Mfap4 | 76293 | -6,03 | 1,88E-02 |
| Abi3bp | 320712 | -6,49 | 2,96E-03 |
| Fmod | 14264 | -6,73 | 1,18E-02 |
| Svep1 | 64817 | -7,27 | 1,92E-02 |
| Islr | 26968 | -7,52 | 9,66E-03 |
| Tpsb2 | 17229 | neg inf | 2,21E-02 |
| Ptx3 | 19288 | neg inf | 4,81E-02 |
| Mmrn1 | 70945 | neg inf | 2,70E-02 |

| | | | |
|---|---|---|---|
| C1qtnf3 | 81799 | neg inf | 1,72E-03 |
| Moxd1 | 59012 | neg inf | 1,85E-03 |
| Olfml2a | 241327 | neg inf | 2,91E-03 |
| Ms4a4d | 66607 | neg inf | 2,05E-02 |
| Adamts16 | 271127 | neg inf | 2,36E-03 |
| Angptl1 | 72713 | neg inf | 3,44E-02 |
| C1qtnf9 | 239126 | neg inf | 2,62E-02 |

**Supplementary Table 5: GSEA identified enriched gene sets**

n=27 (25 enriched in liver, 2 in peritoneal tumors) p-value < 0.01 and FDR < 0.1

| NAME | SIZE | ES | NES | NOM p-val | FDR q-val | FWER p-val | RANK AT MAX |
|---|---|---|---|---|---|---|---|
| GO_POSITIVE_REGULATION_OF_DEFENSE_RESPONSE | 16 | 0,925 | 1,90 | 0,003 | 0,044 | 0,033 | 12 |
| GO_POSITIVE_REGULATION_OF_PHOSPHORUS_METABOLIC_PROCESS | 38 | 0,834 | 1,71 | 0,000 | 0,090 | 0,278 | 12 |
| GO_REGULATION_OF_APOPTOTIC_SIGNALING_PATHWAY | 18 | 0,870 | 1,82 | 0,000 | 0,081 | 0,297 | 12 |
| GO_APOPTOTIC_SIGNALING_PATHWAY | 25 | 0,852 | 1,82 | 0,003 | 0,070 | 0,331 | 12 |
| GO_POSITIVE_REGULATION_OF_PROTEIN_METABOLIC_PROCESS | 51 | 0,773 | 1,82 | 0,000 | 0,063 | 0,335 | 12 |
| GO_POSITIVE_REGULATION_OF_INTRACELLULAR_SIGNAL_TRANSDUCTION | 43 | 0,800 | 1,80 | 0,000 | 0,060 | 0,405 | 12 |
| GO_POSITIVE_REGULATION_OF_PROTEIN_MODIFICATION_PROCESS | 39 | 0,839 | 1,80 | 0,000 | 0,057 | 0,416 | 12 |
| GO_REGULATION_OF_SYSTEM_PROCESS | 26 | 0,825 | 1,80 | 0,006 | 0,061 | 0,458 | 15 |
| GO_SIGNAL_TRANSDUCTION_BY_PROTEIN_PHOSPHORYLATION | 40 | 0,805 | 1,78 | 0,000 | 0,073 | 0,568 | 12 |
| GO_IMMUNE_RESPONSE_REGULATING_SIGNALING_PATHWAY | 17 | 0,888 | 1,78 | 0,006 | 0,071 | 0,583 | 12 |
| GO_REGULATION_OF_MAPK_CASCADE | 36 | 0,814 | 1,78 | 0,000 | 0,074 | 0,615 | 12 |
| GO_POSITIVE_REGULATION_OF_ESTABLISHMENT_OF_PROTEIN_LOCALIZATI | 17 | 0,872 | 1,77 | 0,009 | 0,074 | 0,639 | 12 |
| GO_REGULATION_OF_DEFENSE_RESPONSE | 27 | 0,822 | 1,77 | 0,003 | 0,073 | 0,651 | 12 |
| GO_REGULATION_OF_PROTEIN_MODIFICATION_PROCESS | 66 | 0,741 | 1,77 | 0,000 | 0,076 | 0,694 | 12 |
| GO_NEGATIVE_REGULATION_OF_CELL_DEATH | 35 | 0,789 | 1,76 | 0,006 | 0,085 | 0,747 | 17 |
| GO_POSITIVE_REGULATION_OF_CELL_DEVELOPMENT | 27 | 0,820 | 1,75 | 0,008 | 0,088 | 0,777 | 12 |
| GO_PLATELET_ACTIVATION | 20 | 0,862 | 1,74 | 0,006 | 0,089 | 0,805 | 12 |
| GO_POST_TRANSLATIONAL_PROTEIN_MODIFICATION | 19 | 0,870 | 1,74 | 0,006 | 0,087 | 0,807 | 9 |
| GO_NEGATIVE_REGULATION_OF_RESPONSE_TO_EXTERNAL_STIMULUS | 25 | 0,827 | 1,73 | 0,006 | 0,093 | 0,857 | 12 |
| GO_APOPTOTIC_PROCESS | 68 | 0,709 | 1,73 | 0,000 | 0,093 | 0,868 | 35 |
| GO_PEPTIDE_HORMONE_SECRETION | 17 | 0,863 | 1,72 | 0,009 | 0,094 | 0,913 | 15 |
| GO_REGULATION_OF_PROTEIN_LOCALIZATION | 37 | 0,765 | 1,72 | 0,003 | 0,087 | 0,916 | 15 |
| GO_REGULATION_OF_PHOSPHORUS_METABOLIC_PROCESS | 67 | 0,728 | 1,71 | 0,000 | 0,091 | 0,934 | 12 |
| GO_PROTEIN_PHOSPHORYLATION | 68 | 0,729 | 1,71 | 0,000 | 0,089 | 0,934 | 12 |
| GO_REGULATION_OF_BODY_FLUID_LEVELS | 39 | 0,769 | 1,71 | 0,008 | 0,090 | 0,94 | 27 |
| GO_EXTRACELLULAR_MATRIX_COMPONENT | 22 | -0,817 | -1,84 | 0,000 | 0,016 | 0,02 | 87 |
| GO_COLLAGEN_TRIMER | 16 | -0,862 | -1,81 | 0,000 | 0,023 | 0,055 | 87 |
